# Microswitch-Guided Sampling for the Detection of Ligand Signaling Bias in GPCR Systems

**DOI:** 10.64898/2026.09.16.752090

**Authors:** Paulina Dragan, Dorota Latek

## Abstract

Biased signaling occurs when a ligand preferentially activates a specific signaling pathway at a certain receptor within a particular cellular environment. As pharmaceuticals, functionally selective (biased) compounds can engage beneficial signaling pathways while avoiding those linked to adverse effects, resulting in safer and targeted therapeutics. However, existing experimental methods for characterizing ligand bias are often time-consuming and costly, and they hardly elucidate the structural basis of biased signaling. Here, we present microswitch-guided sampling (MGS), a computational method that uses conformational changes in molecular switches to assess ligand bias in G protein-coupled receptor systems. Using initial 2 µs all-atom molecular dynamics simulations of the β_1_-adrenergic receptor (β_1_AR), the chemokine receptor CXCR3, and the µ-opioid receptor (µOR) in complex with either G protein or β-arrestin and known biased ligands, we identified microswitches whose conformational changes were linked to activation of a specific signaling pathway. We then sampled simulation trajectories for representative complex conformations selected at these microswitch-change time points and used them as starting points for 500 ns ligand-swapped simulations, in which we exchanged biased ligands between complexes. We observed clear differences in conformational changes between complexes bound to pathway-activating and non-activating ligands in at least two of three replicas across all examined systems, demonstrating that MGS can distinguish G protein-biased from β-arrestin-biased ligands. These findings show that MGS is an effective tool for assessing signaling pathway bias in GPCR systems and suggest that this methodology can be applied to other GPCR targets to act as a fast, ns-scale computational assay to detect functional selectivity with molecular dynamics.

**Significance statement:** Biased ligands that activate specific signaling pathways at G protein-coupled receptors hold considerable promise as safer therapeutics, but identifying and characterizing them remains experimentally challenging. Here, we present microswitch-guided sampling (MGS), a computational method that uses conformational changes in conserved molecular switches to assess ligand bias in GPCR–transducer complexes via all-atom molecular dynamics simulations. By identifying pathway-specific conformational changes and testing them across ligand-swapped complexes, MGS distinguished G protein-biased from β-arrestin-biased ligands across distinct GPCR systems. This establishes MGS as a time- and cost-effective computational alternative to functional assays for characterizing ligand signaling bias, to accelerate the design of functionally selective drugs for GPCR targets.

## 1. Introduction

Biased ligands, acting at a certain G protein-coupled receptor (GPCR) within a specific cellular environment, preferentially activate one signaling pathway over another. This occurs because biased ligands can stabilize different active receptor conformations, allowing more selective binding to a single transducer (1). Canonically, biased agonism is described in terms of two signaling pathways—G protein (2) or β-arrestin-mediated (3) (**Figure 1**). However, the biased ligand alone does not determine pathway-specific bias. System bias, arising from differences, e.g., in the expression levels of receptors, transducers, effectors, or modulatory proteins, can also affect a receptor’s signaling profile. The signaling outcome resulting from both ligand and system bias is termed functional selectivity (1).

**Figure 1.**
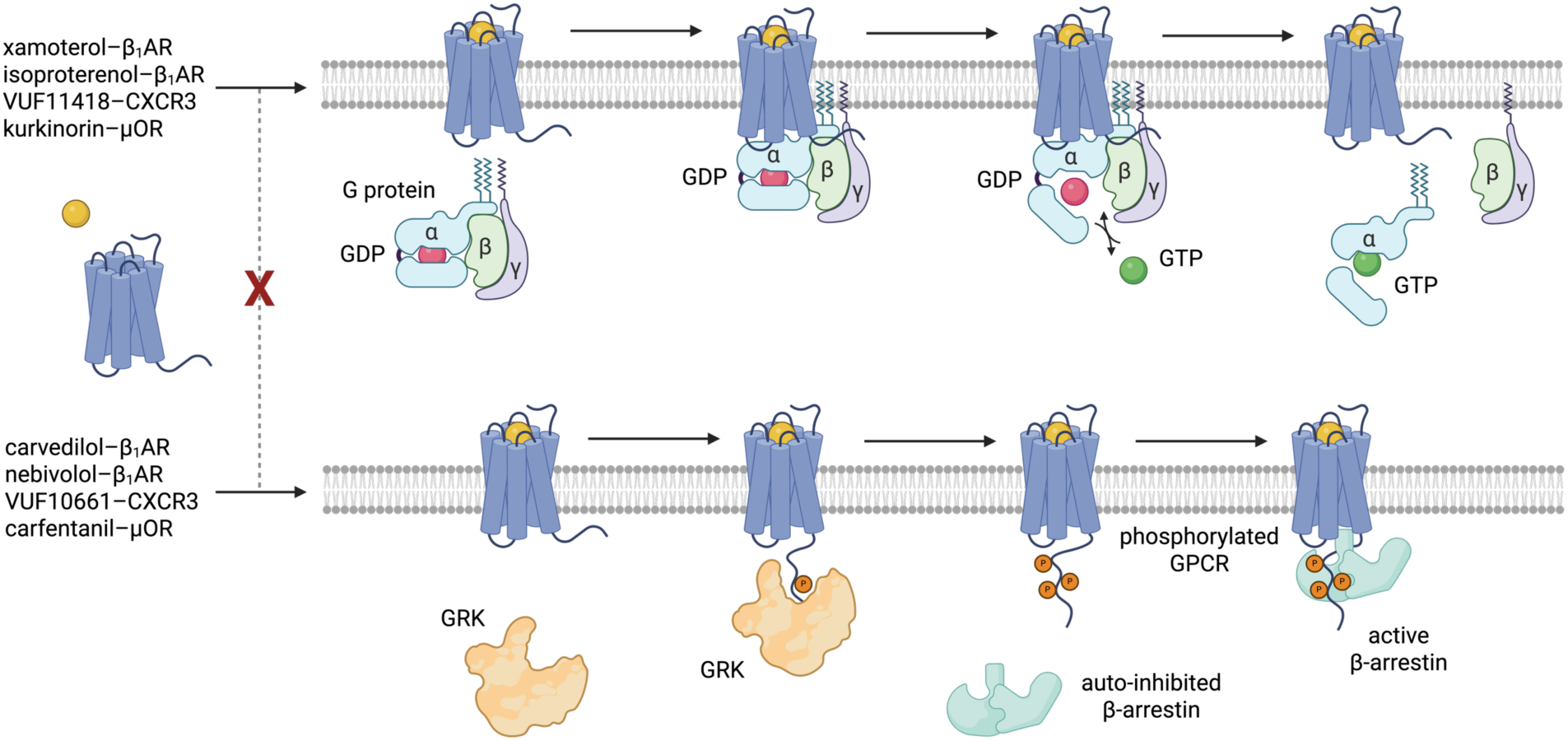
Ligand-induced functional selectivity in GPCR systems. (Top) A schematic representation of G protein activation: complex formation, GDP/GTP exchange, and G protein dissociation. **(Bottom)** A schematic representation of β-arrestin activation: GRK recruitment, receptor phosphorylation, and β-arrestin binding. Created using BioRender.

Controlling signaling bias has significant therapeutic implications. Since downstream signaling pathways mediate distinct physiological effects, it is possible to design drugs that activate only a therapeutically beneficial pathway while avoiding others associated with adverse effects. This results in safer, more efficient drugs with high specificity for disease-affected tissue, e.g., a targeted therapy for cancer (4). Certain β-blockers target β-adrenergic receptors and promote cardioprotection by activating the β-arrestin signaling pathway while simultaneously inhibiting G protein activation (5, 6). This, along with the opioid receptor system, has prompted research to identify and/or design biased ligands for other GPCR targets.

In experimental studies, ligand signaling bias can be determined by comparing a ligand’s ability to activate a specific signaling pathway with that of an unbiased reference ligand using specific functional assays (1, 7). While reliable, this approach is time-consuming and expensive because it requires multiple assays to determine a ligand bias (8). Additionally, functional assays do not directly reveal the structural basis of the observed functional selectivity (8). In contrast, determining the structure of each biased ligand complex using standard X-ray crystallography and cryo-EM is costly and time-consuming. Moreover, these static methods provide snapshots of possible receptor conformations, offering limited insight into the detailed step-by-step mechanisms of GPCR and transducer activation. However, more advanced experimental methods, such as XFEL or time-resolved cryo-EM, are being developed to capture protein dynamics (9). In addition, computational methods such as molecular dynamics (MD) enable simulation of GPCR receptors bound to biased ligands and transducers, capturing atom-level conformational changes (9). These include changes in micro- and macroswitches (10), in which evolutionarily conserved residues undergo conformational changes to guide protein activation and signaling via a particular pathway (11, 12), as exemplified by our previous study of the orexin receptor system (13, 14) and studies performed by others (15, 16). Based on these preliminary studies, we assumed that alterations in microswitch conformations might indicate not only receptor activation but also ligand-induced signaling bias.

In the current study, we examined trajectories from 2 µs-long, explicit membrane, all-atom MD simulations of the β_1_ adrenergic receptor (β_1_AR, encoded by the ADRB1 gene), the chemokine receptor CXCR3 (encoded by the CXCR3 gene), and the µ-opioid receptor (µOR, named also MOR-1, encoded by the OPRM1 gene) in complex with biased ligands and different transducers. We identified time points when conformations of certain microswitches and macroswitches in transducers changed, indicating system activation. We then sampled the simulation trajectory across frames adjacent to these time points to identify representative conformations of the ligand– GPCR–transducer complex just before the activation-like change of the system. Such an approach formed the basis of the microswitch-guided sampling (MGS) method developed and described here. Next, to determine if a particular microswitch was exclusively linked to activation of a specific signaling pathway, rather than responding to non-specific agonist binding and unbiased receptor/transducer activation, we interchanged the biased ligands between complexes. Molecular switches that consistently changed (or failed to change) their conformations according to the considered signaling-pathway bias were chosen as pathway-specific markers of biased signaling. Representative ligand–GPCR–transducer complex conformations selected from simulations based on microswitch change time points were used in a computational assay to assess the signaling bias of tested ligands. In the following GPU-accelerated, 500 ns-long, explicit membrane, all-atom MD simulations, we observed clear differences in conformational changes of transducers upon differently biased ligands enabling their time- and cost-saving identification in terms of the signaling bias. Since activation-induced conformational changes in transducers are similar across many (though not all) GPCR systems (17), the microswitches identified here as pathway-specific markers may be used to develop similar computational assays to test biased agonism of other GPCR ligands.

## 2. Results

### 2.1. G Protein Complexes

#### 2.1.1. β_1_AR–Gs

Initial simulations of β_1_AR–Gs complexes were performed with the G protein-biased ligand xamoterol (5). Analysis of the initial simulations showed the most distinct conformational changes in switches II and III, suggesting a transition state, as evidenced by an increase in the distance between Cζ of R228^G.H2.01^ and Cδ of E259^G.4h3.10^ (**Figure 2A**). This conformational change in switches II and III underlies MGS in β_1_AR–Gs complexes. From each replica, the frame 10 ns before the most significant change (here, the largest increase in distance) was exported. In the ligand-swapped simulations, where xamoterol was replaced with the β-arrestin-biased carvedilol (5), an increase in the distance between Cζ of R228^G.H2.01^ and Cδ of E259^G.4h3.10^ was not observed. However, the differences between the xamoterol and carvedilol replicas were not clear enough to use this switch alone to draw conclusions about the ligand bias. As such, heavy-atom RMSD values for both the entire Gα subunit and the α-helical domain (AHD) were computed instead (**Figure 2B**). To make the initial and ligand-swapped simulations comparable, RMSD was computed over 500 frames in each replica, starting from the frame exported for MGS. Gα showed higher RMSD values in the initial simulations than in the ligand-swapped simulations, indicating larger-scale conformational changes. The RMSD differences for AHD were less evident, though they had a higher baseline in the xamoterol replicas. However, because the comparison spans only 500 ns, it may not have been sufficient to clearly distinguish AHD separation from the Ras-like domain. On the other hand, RMSD fluctuations across the entire Gα subunit could suggest activation-like conformational changes beyond AHD.

**Figure 2.**
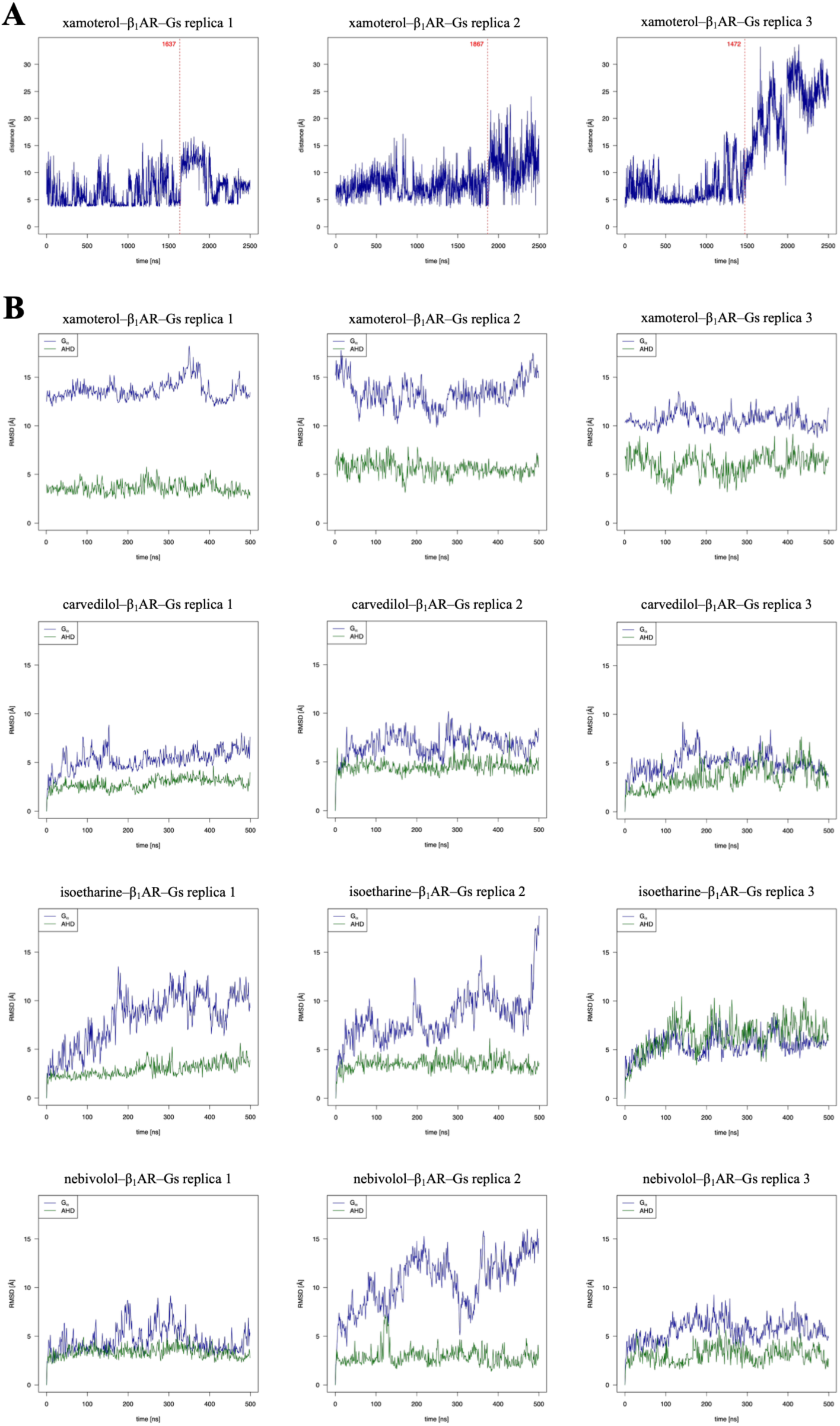
**(A)** The distance between Cζ of R228^G.H2.01^ and Cδ of E259^G.4h3.10^ in the xamoterol–β_1_AR–Gs complexes. The frames exported for MGS are indicated in red. **(B)** The heavy-atom RMSD of G*a* (superposed receptors) and the helical domain (superposed Ras-like domain) for the initial (xamoterol), ligand-swapped (carvedilol), and test (isoetharine and nebivolol) β_1_AR–Gs complexes. For the xamoterol replicas, RMSD was computed starting from the same frame exported for MGS. The simulation trajectory analysis was limited to the first 500 ns.

To assess whether these results were signaling-bias specific, we performed a third set of simulations with two ligands: isoetharine, reported as preferentially activating β-arrestin-mediated signaling in β_2_AR (18) but exhibiting G protein signaling bias in β_1_AR (19), and nebivolol—a β-arrestin-biased ligand in both β_1_AR and β_2_AR (5). All three replicas of the isoetharine-bound complex showed indications of continued G protein activation, consistent with its bias toward the G protein signaling pathway in β_1_AR (19) (**Figure 2B**). This is especially evident in replicas 1 and 2, where Gα RMSD began to increase at the start of the simulation, though it reached values slightly lower than those for xamoterol. Gα RMSD was lower in replica 3, but AHD RMSD increased to values not observed for either β-arrestin-biased ligand, suggesting G protein bias for isoetharine. In contrast, replicas 1 and 3 of the nebivolol-bound complexes demonstrated low RMSD values for both the entire Gα and the AHD, suggesting an inhibition of G protein activation. The increased Gα RMSD observed for replica 2 reflects non-activation-related conformational fluctuations within Gα, given the simultaneous low AHD RMSD. As shown in **Figure 12Error! Reference source not found.** (red line), simulation systems with an oppositely biased ligand may fluctuate around a local minimum without demonstrating further activation-like or deactivation-like conformational changes. Both behaviors, either fluctuating or deactivation-like, differ significantly from the activation-like conformational changes of the reference system and thus represent markers for detecting the oppositely biased ligand in the computational assay described here.

These results are broadly consistent with a PCA of each trajectory (**Figure 3**, **Supplementary Figure S1**). In the xamoterol-bound replicas, early and late frames form distinct clusters across multiple principal components (PCs), with multimodal histograms indicating sampling of multiple conformational states. Replica 1 shows the clearest state separation in the PC1/PC2 projection. In the carvedilol-including replicas, time ordering is less consistent, and the scatter distributions are more diffuse, particularly in replicas 1 and 3. The intermixing of early and late frames is consistent with reduced conformational change of the carvedilol-including replicas relative to xamoterol. For the isoetharine and nebivolol systems, PCA results from 500 ns simulations are slightly less readable because of the much shorter simulation time (fewer frames), compared with 2 µs for the reference ligand simulations or 1 µs for the ligand-swapped simulations. However, they are still sufficient to draw reliable conclusions from the PCA landscape—the streak-like distributions indicate that simulation systems have not moved from their starting conformation to a significant extent but rather fluctuate locally. Nevertheless, the nebivolol-bound replicas show greater overlap between early and late frames than the isoetharine replicas, which is qualitatively consistent with the RMSD analysis showing no signs of activation-like changes in Gα in replicas 1 and 3, including nebivolol. These observations support the possibility of evaluating both ligands in terms of their varied functional selectivity. Notably, the consistency between the RMSD results and PCA confirms that PCA could strongly support the interpretation of the computational assay results. Both the PCA and RMSD analyses showed that G protein-biased ligands promote larger-scale conformational rearrangements in β_1_AR–Gs complexes than the oppositely biased ligands.

**Figure 3.**
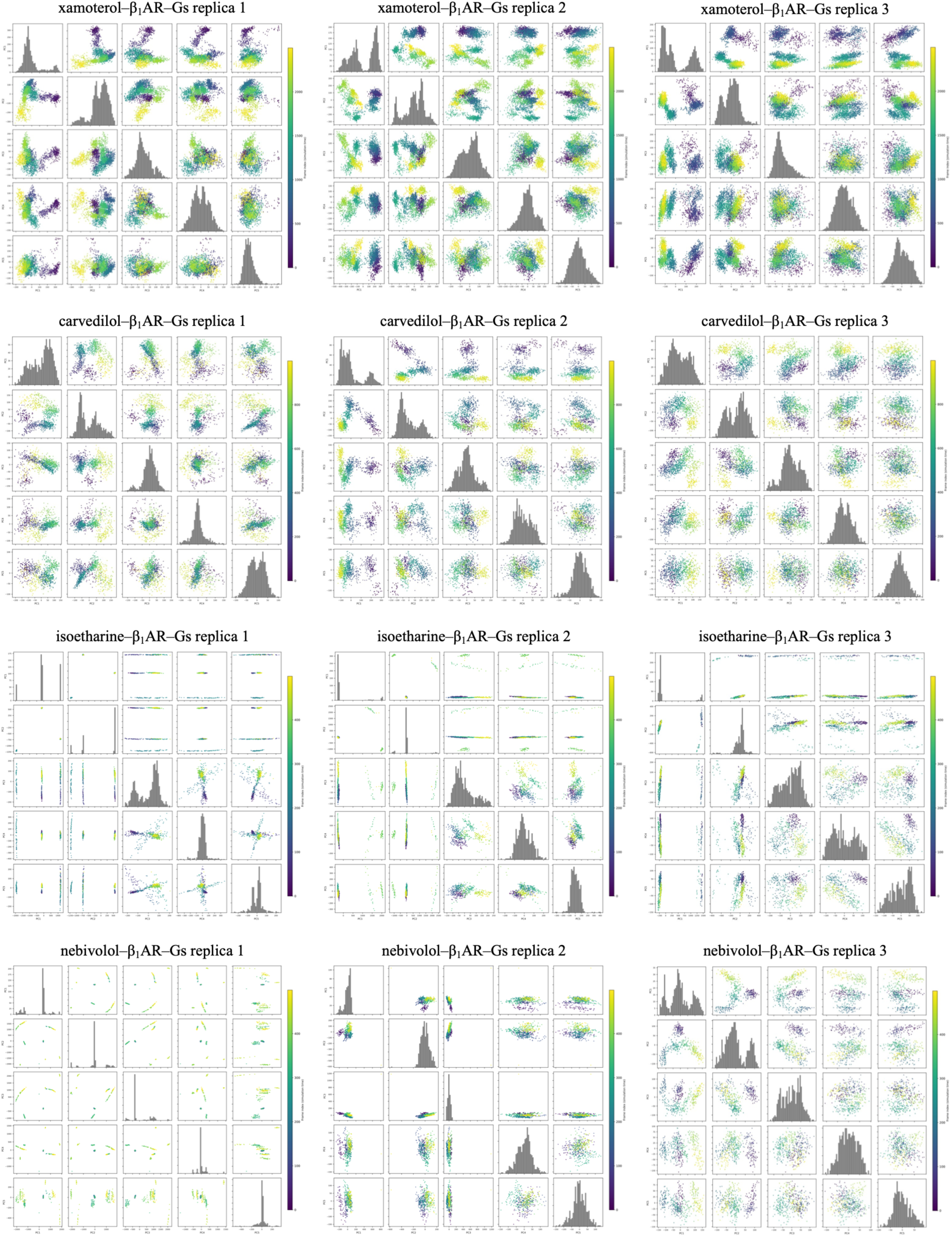
Principal component analysis of the β_1_AR–Gs simulations. Pairwise scatter plots of the projections of the trajectory onto the first 5 PCs, colored by frame index (simulation time; purple = early, yellow = late). Diagonal panels show the distribution of each PC. The analyses were performed on protein backbone atoms following structural alignment. Scatter plots of the first 10 PCs are provided in Supplementary Figure S6.

#### 2.1.2. CXCR3–Gi

Simulations of CXCR3–Gi complexes were performed with the G protein-biased VUF11418 (20) and the β-arrestin-biased VUF10661 (20). D295^G.HG.02^ was selected as the microswitch used to identify frames for MGS, as it exhibits a clear change in the χ1 dihedral angle (N–C_α_–C_β_–C_γ_) upon shift from an inactive to an active conformation (∼60° to − 60°, respectively) as observed in our preliminary study of Gq activation by a non-canonical biased ligand in the orexin receptor system (13, 14). As with β_1_AR, for each replica, the exported frame was taken 10 ns before the microswitch changed from an inactive-like to an active-like conformation. However, in the ligand-swapped simulations, where VUF11418 was replaced with the β-arrestin-biased VUF10661, D295^G.HG.02^ did not always remain in an inactive state as expected; this was also observed for the reference, initial simulations with VUF11418 (**Figure 4**). This suggests that D295^G.HG.02^ in the CXCR3 systems tends to fluctuate to a certain point regardless of the ligand type. Accordingly, conformational changes in the remaining microswitches were examined to identify observable differences between the initial (reference) and ligand-swapped simulations. The most evident differences were observed in microswitches II and III. As in the β_1_AR–Gs complexes, G protein activation increased the distance between them, as evidenced by the breakage of interactions between R205^G.H2.01^ and E236^G.4h3.10^. To make the initial and ligand-swapped simulations comparable, we computed hydrogen bond formation between these two residues over 500 frames in each replica, starting from the frame exported for MGS (**Figure 4A**). This hydrogen bond was absent in simulations with the G protein-biased ligand but present in those with the β-arrestin-biased ligand, confirming it could indicate ligand signaling bias in the CXCR3 system (**Figure 4B**).

**Figure 4.**
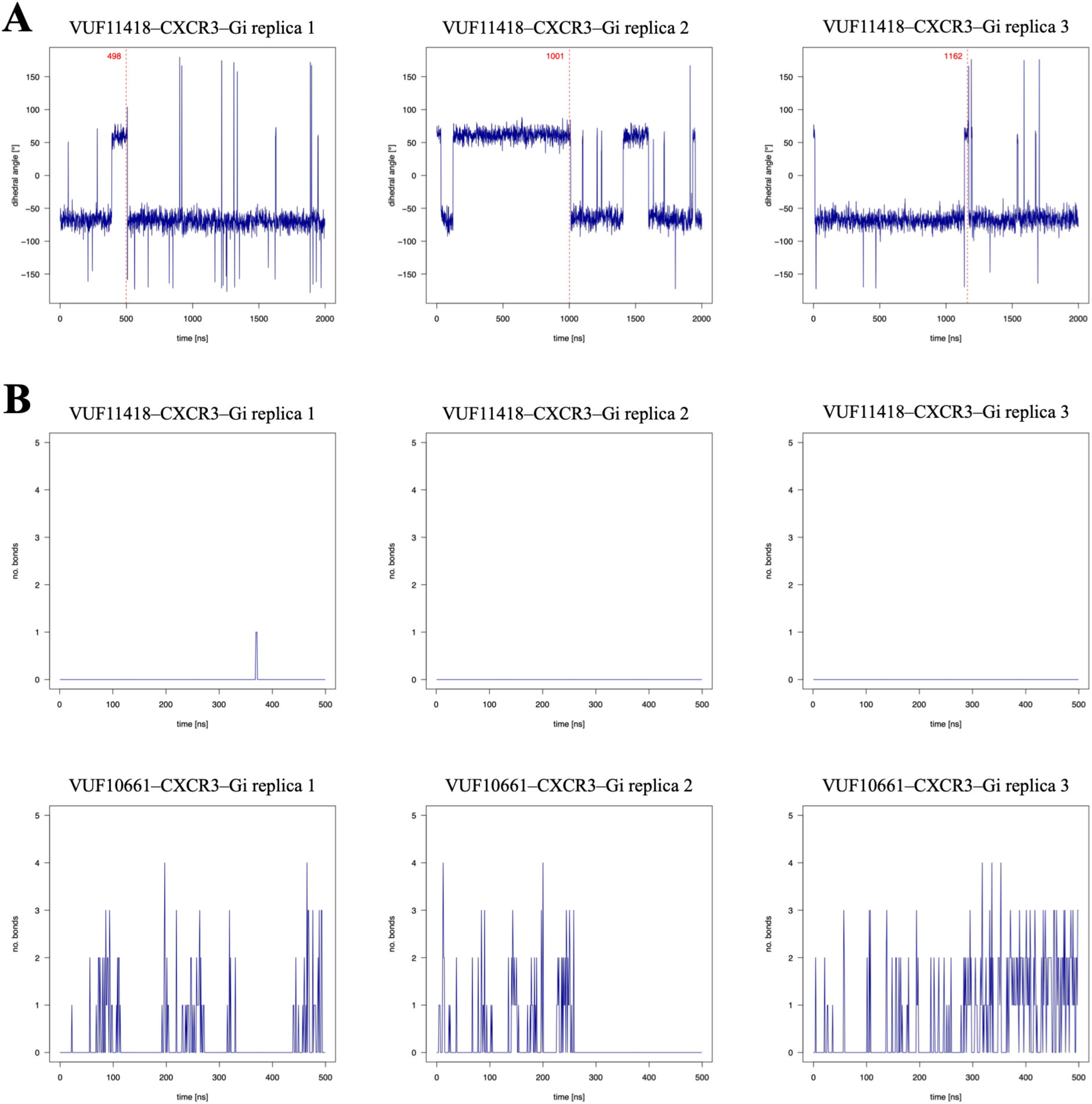
**(A)** Changes in the χ1 dihedral angle (N–C_α_–C_β_–C_γ_) of D295^G.HG.02^ in the VUF11418–CXCR3–Gi complexes. The frames exported for use with MGS are indicated in red. **(B)** Formation of hydrogen bonds between R205^G.H2.01^ and E236^G.4h3.10^ in CXCR3–Gi complexes with the G protein-biased ligand (VUF11418) and β-arrestin-biased ligand (VUF10661). For the VUF11418 replicas, hydrogen bond formation was computed starting from the same frame exported for MGS. The simulation trajectory analysis was limited to the first 500 ns.

The PCA results (**Figure 5**, **Supplementary Figure S2**) largely agree with the hydrogen-bond analysis above. Among the VUF11418-bound replicas, replicas 1 and 2 show the most clearly structured PC1/PC2 projections, with the transition from early to late frames forming an arc. As with the xamoterol–β_1_AR–Gs replicas, it indicates the appearance of multiple distinct states and ongoing conformational changes resulting in G protein activation. In contrast, the data points for replica 3 are more diffuse and more closely resemble the landscape observed for the VUF10661 replicas, suggesting this simulation is slightly less consistent with G protein activation, possibly because it traps the system in a local minimum. This again confirms the need for multiple-replica simulations in the described computational assay due to the limitations of conventional MD. In the VUF10661-bound replicas, data point distributions are generally diffuse, and early- and late-frame data points are intermixed in replicas 1 and 3, suggesting system fluctuations rather than ongoing, activation-like conformational changes. The more structured behavior of replica 2 is consistent with the results presented in **Figure 4B**, where hydrogen bonds between R205^G.H2.01^ (switch II) and E236^G.4h3.10^ (switch III) broke partway through the simulation. This may result from VUF10661, like VUF11418, being partially rather than fully biased toward β-arrestin and Gi, respectively (20). However, together with the hydrogen-bond results, the PCA results confirmed that VUF11418 more consistently induces activation-like conformational changes in Gi than VUF10661, with inter-replica variability reflecting partially overlapping signaling profiles due to partial rather than fully biased agonism.

**Figure 5.**
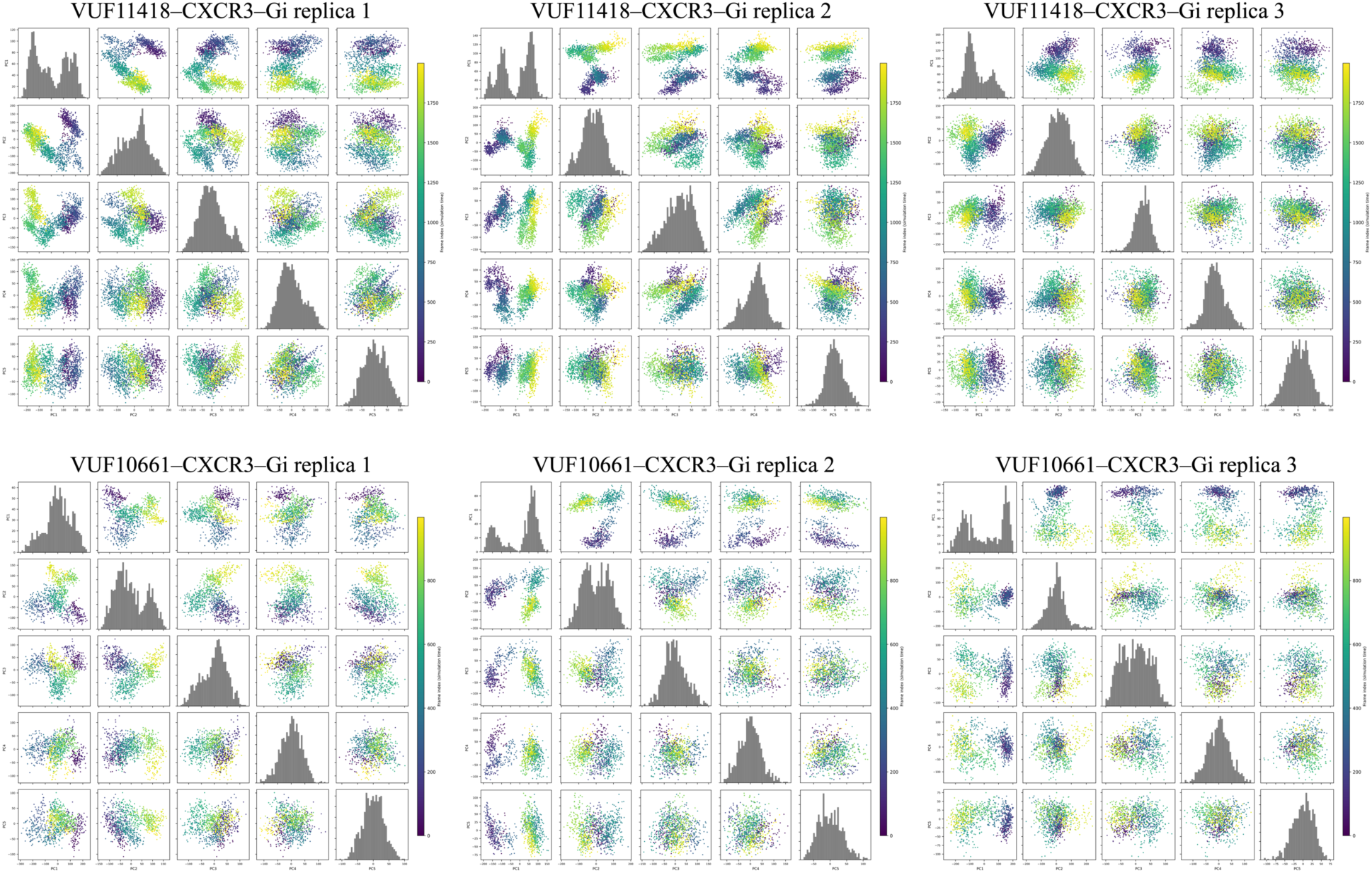
Principal component analysis of the CXCR3–Gi simulations. Pairwise scatter plots of the projections of the trajectory onto the first 5 PCs, colored by frame index (simulation time; purple = early, yellow = late). Diagonal panels show the distribution of each PC. The analyses were performed on protein backbone atoms following structural alignment. Scatter plots of the first 10 PCs are provided in Supplementary Figure S7.

#### 2.1.3. µOR–Gi

Simulations of the µOR–Gi complexes were performed with the G protein-biased kurkinorin (21) and the β-arrestin-biased carfentanil (22). As with the xamoterol–β_1_AR-Gs replicas, the distance between microswitches II and III was used to identify frames for MGS (**Figure 6A**). For each trajectory, the exported frame was taken before a sharp increase in the distance between the G*a* residues R205^G.H2.01^ and E236^G.4h3.10^. As these changes were not clearly observable in the ligand-swapped simulations, hydrogen bond formation between R205^G.H2.01^ and E236^G.4h3.10^ was examined instead (**Figure 6B**). Again, while hydrogen bonds were largely absent in the initial (reference) replicas, they formed in 2 of the 3 ligand-swapped replicas, further confirming the usefulness of interactions between these two residues for assessing ligand signaling bias.

**Figure 6.**
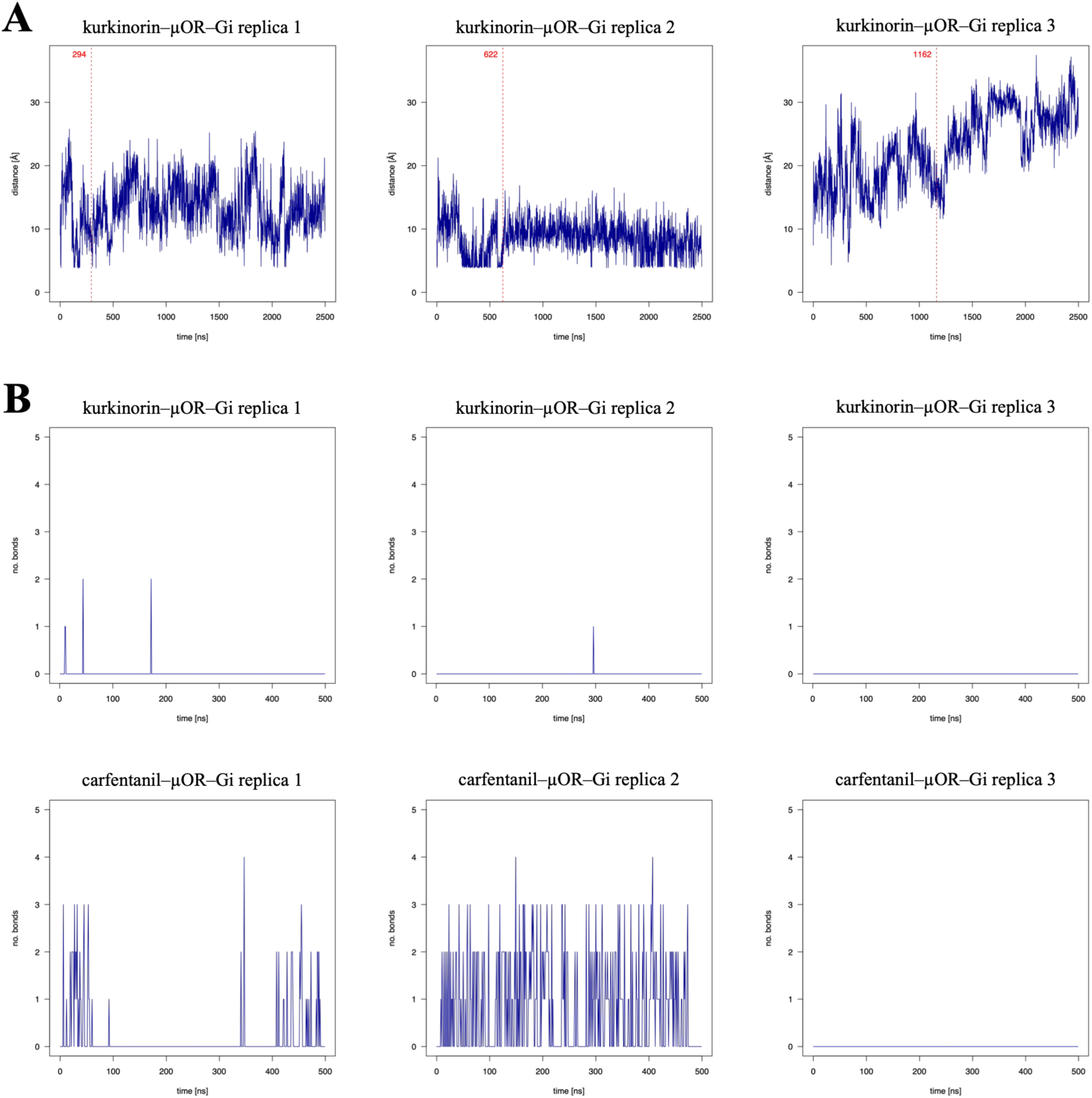
**(A)** The distance between Cζ of R205^G.H2.01^ and Cδ of E236^G.4h3.10^ in the kurkinorin–µOR–Gi complexes. The frames exported for use with MGS are indicated in red. **(B)** Formation of hydrogen bonds between R205^G.H2.01^ and E236^G.4h3.10^ in µOR–Gi complexes with the G protein-biased ligand (kurkinorin) and β-arrestin-biased ligand (carfentanil). For the kurkinorin replicas, hydrogen bond formation was computed starting from the same frame exported for MGS. The simulation trajectory analysis was limited to the first 500 ns.

Again, the PCA results (**Figure 7**, **Supplementary Figure S3**) were broadly consistent with the hydrogen-bond analysis above for the kurkinorin-bound replicas, though slightly less so for the carfentanil-bound complexes. All three kurkinorin-bound replicas demonstrated clearly structured PC1/PC2 projections with strong time-ordering, multimodal histograms, and scree plots confirming that a single dominant mode accounts for most of the variance. This suggests conformational changes consistent with G protein activation, especially given the lack of hydrogen bonds in **Figure 6B**. The replica 3 trajectory shows a pronounced sweeping progression in PC1/PC2, suggesting it underwent the most extensive conformational change of the three. The scree plot, with PC1 exceeding 40%, confirms this as a single dominant motion. Surprisingly, the carfentanil-bound replicas also show time-ordering. In replica 1, multiple histogram peaks and a flatter scree plot suggest a less directed conformational change. Replicas 2 and 3 show more structured arcs in PC1/PC2, and their scree plots are more comparable to those obtained for the kurkinorin complexes. However, given that hydrogen bonds between R205^G.H2.01^ and E236^G.4h3.10^ were present in replica 2, the directional conformational change visible in the PCA for this replica reflects a conformational change that is not associated with G protein activation, as in the case of replica 2 in the nebivolol–β_1_AR–Gs system (see above). Longer simulation of minimum 2 µs, considering results of the initial simulations, could reveal if these conformational changes indeed lead to the inactive state I (Figure 2). However, it was not the aim of this study, because to test many ligands in a computational assay in a relatively short time, the maximum simulation time had to be limited to 500 ns.

**Figure 7.**
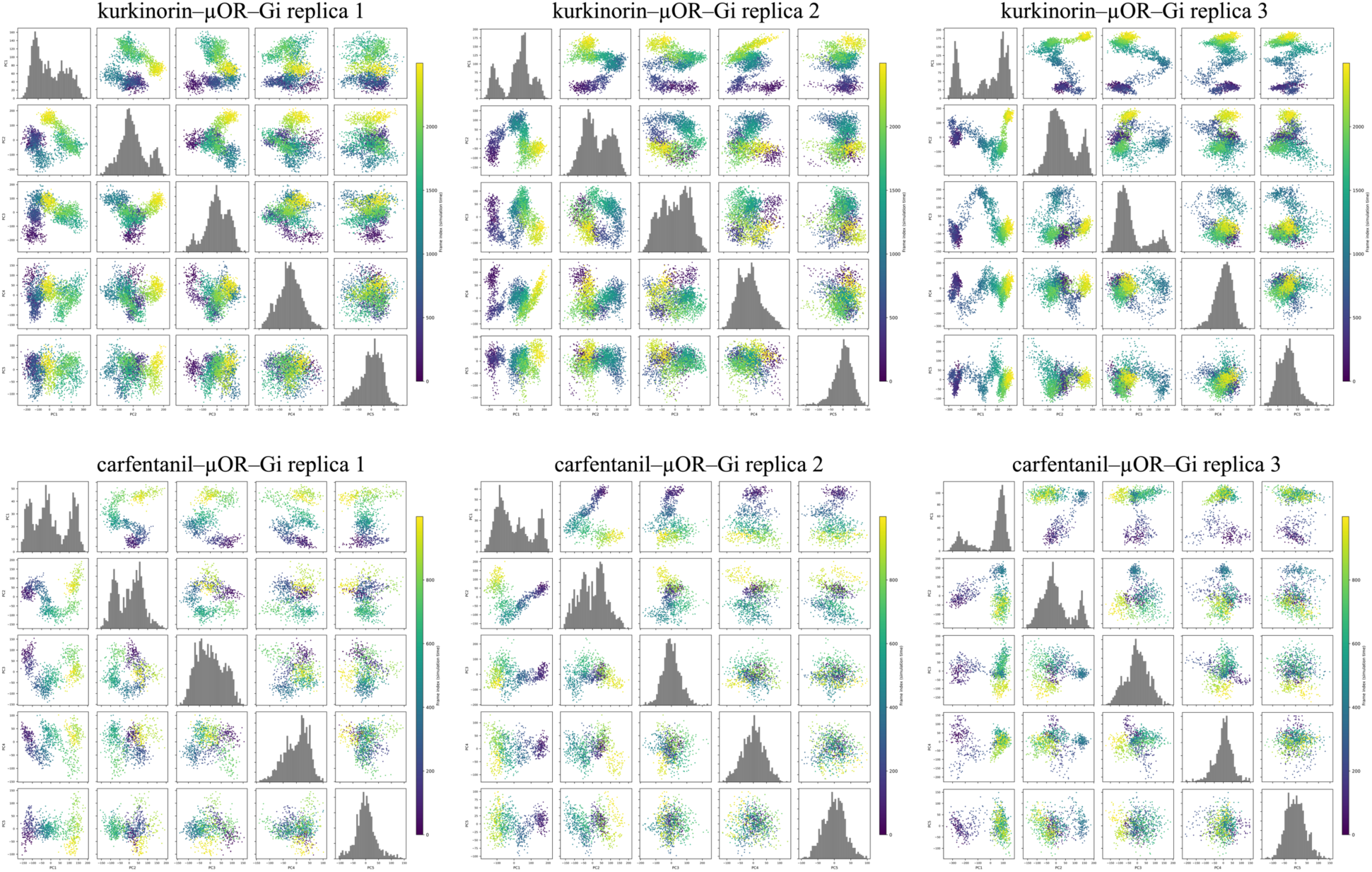
Principal component analysis of the µOR–Gi simulations. Pairwise scatter plots of the projections of the trajectory onto the first 5 PCs, colored by frame index (simulation time; purple = early, yellow = late). Diagonal panels show the distribution of each PC. The analyses were performed on protein backbone atoms following structural alignment. Scatter plots of the first 10 PCs are provided in Supplementary Figure S8.

### 2.2. β-Arrestin Complexes

#### 2.2.1. β_1_AR–β-arrestin

Initial simulations of β_1_AR–β-arrestin complexes included the β-arrestin-biased ligand carvedilol. The most noticeable differences among replicas were observed in the RMSD of the β-arrestin finger loop (**Figure 8A**). The corresponding frames 10 ns before the most significant RMSD change were exported for MGS. The finger-loop RMSD was also examined in the ligand-swapped and ligand-test simulations, though these results were less conclusive. Accordingly, conformational changes in the finger and middle loops of β-arrestin were also examined and overlaid on the RMSD plots (**Figure 8B**). Again, for comparability, the RMSD for the initial simulations was computed over the first 500 ns in each replica, starting from the frame exported for MGS. The carvedilol-bound replicas exhibited the highest RMSD values across all β_1_AR–β-arrestin complexes. In all three replicas, the finger loop began to lift upward to enter the receptor, indicating the ongoing β-arrestin binding to β_1_AR and its subsequent activation. In contrast, the xamoterol-bound complexes had much lower RMSD values, and the finger loop conformation changed less. Although the ligand-swapped simulations were shorter than the initial simulations (1 µs vs. 2 µs), the timescale was long enough to observe clear changes in the finger-loop conformation if β-arrestin activation had progressed. Instead, especially in xamoterol-bound replicas 1 and 3, the finger loop remained mostly in the same conformation in the first and final frames, as in the middle loop of replica 1. While replica 2 showed slightly larger conformational fluctuations, they were still too small to account for continued β-arrestin activation. In the ligand-test simulations, the conformational changes were larger. In particular, in the nebivolol-bound replicas, the finger loop continued to demonstrate activation-like conformational changes, lifting upward relative to the starting conformation in two of three replicas. In contrast, the conformational changes observed in the isoetharine-bound replicas were smaller. This was particularly evident for replicas 2 and 3, where RMSD values were lower than those observed for the corresponding nebivolol-bound replicas. In isoetharine-bound replica 3, RMSD values were larger, but the final middle-loop conformation remained largely similar to the initial conformation exported for MGS.

**Figure 8.**
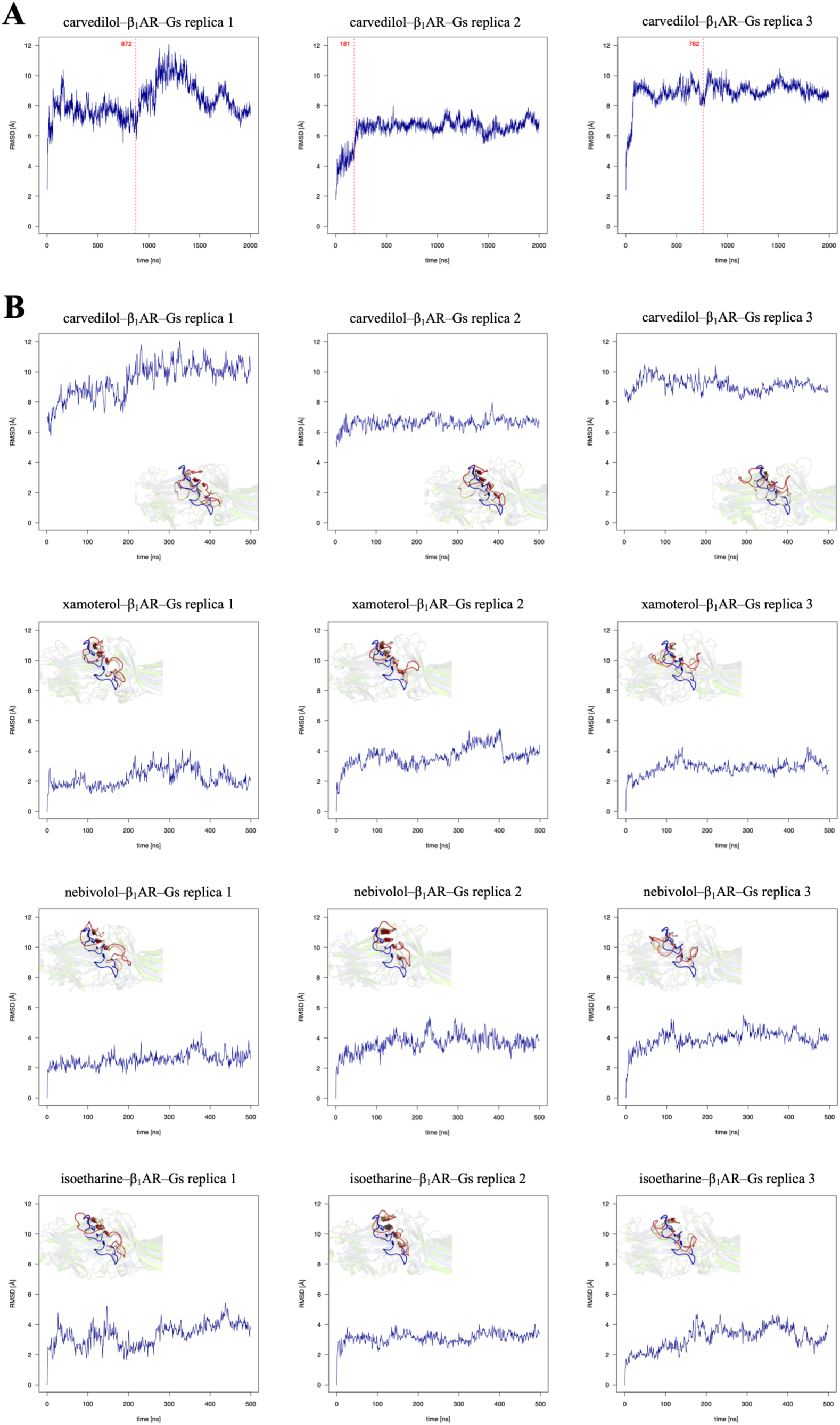
**(A)** The heavy-atom RMSD of the finger loop (superposed receptor) in the carvedilol–β_1_AR–β-arrestin complexes. The frames exported for use with MGS are indicated in red. **(B)** The heavy-atom RMSD of the finger loop (superposed receptor) for the initial (carvedilol), ligand-swapped (xamoterol), and test (nebivolol and isoetharine) β_1_AR–β-arrestin complexes. For the carvedilol replicas, the RMSD was computed starting from the frame exported for MGS. Overlaid are the conformations of the finger and middle loops in the active 6TKO PDB structure (blue), and the first (beige) and last (red) frame of each simulation. The simulation trajectory analysis was limited to the first 500 ns.

The PCA results (**Figure 9**, **Supplementary Figure S4**) resembled the pattern observed in the β_1_AR–Gs complexes, with the ligand-bound complex with its cognate transducer showing the most directed conformational sampling. All the carvedilol-bound replicas show the structured PC1/PC2 projections. In replica 1, distinct clusters instead of a continuous arc suggest that, rather than a gradual transition between conformational changes, the system switched more abruptly from one conformational state to another. In contrast, the conformational landscape in replica 2 is more diffuse, but the transition between early and late frames is more gradual than in replica 1. Replica 3 displays the smoothest transition, with a clear arc shape in the PC1/PC2 projection. In contrast, the xamoterol-bound replicas show more diffuse data points, especially in replica 1. In the same replica, the histogram is multimodal but has less-separated peaks, suggesting less structured conformational sampling. Replica 2 is slightly more structured, but time ordering, especially for higher PCA components, is more diffuse than in the carvedilol-bound complexes. Replica 3 shows clear time ordering and an arc-shaped transition between early and late frames but only for lower components. However, higher-order projections are more diffuse than those in the carvedilol-bound complexes, suggesting conformational fluctuations rather than direct β-arrestin activation. As with the β_1_AR–Gs complexes, the number of frames extracted from the isoetharine and nebivolol simulations is slightly too small to differentiate between isoetharine and nebivolol in terms of β-arrestin-mediated signaling. However, the data point clusters for the isoetharine replicas are more diffuse, suggesting a broader exploration of the conformational landscape than in the nebivolol replicas, indicating transducer fluctuations rather than its activation, respectively.

**Figure 9.**
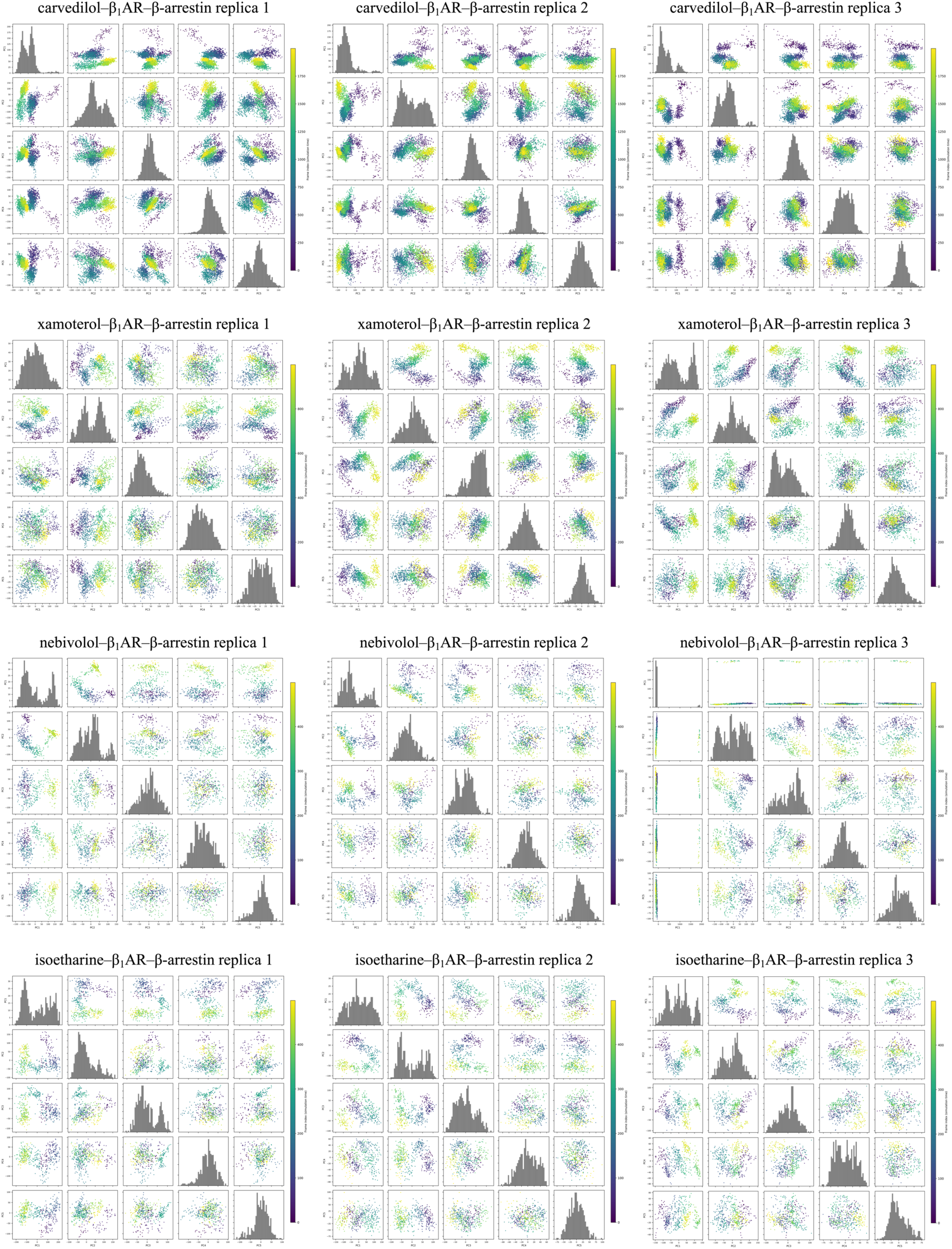
Principal component analysis of the β_1_AR–β-arrestin simulations. Pairwise scatter plots of the projections of the trajectory onto the first 5 PCs, colored by frame index (simulation time; purple = early, yellow = late). Diagonal panels show the distribution of each PC. The analyses were performed on protein backbone atoms following structural alignment. Scatter plots of the first 10 PCs are provided in Supplementary Figure S9.

#### 2.2.2. µOR–β-arrestin

Initial simulations of the µOR–β-arrestin complexes included the β-arrestin-biased ligand carfentanil. Noticeable differences were observed in the distance between residue D69^N.s5s6.06^ of the β-arrestin finger loop and R76^N.S6.02^ in a nearby β-strand. A hydrogen bond between these two residues, together with a hydrogen bond between D67^N.s5s6.04^ and R62^N.S5.11^, forms an ionic lock that keeps the finger loop in an inactive conformation (23). β-arrestin activation breaks this hydrogen bond (23), increasing the distance between these two residues. For each trajectory, the exported frame was taken before a sharp increase in this distance (**Figure 10A**). These changes were less comparable in the subsequent ligand-swapped simulations; instead, RMSD and finger and middle loop conformations were examined, as in the β_1_AR–β-arrestin replicas (**Figure 10B**). Again, the carfentanil-bound replicas demonstrated larger RMSD values and fluctuated more than the kurkinorin-bound replicas in terms of β-arrestin. Finger loop conformations in specific replicas were compared to that in the 9WSV PDB structure (24). In the carfentanil replicas, the finger loops showed clear upward movements relative to the MGS starting conformations (**Figure 10B**). In contrast, finger-loop conformational changes were less pronounced in the kurkinorin replicas, suggesting inhibition of β-arrestin activation.

**Figure 10.**
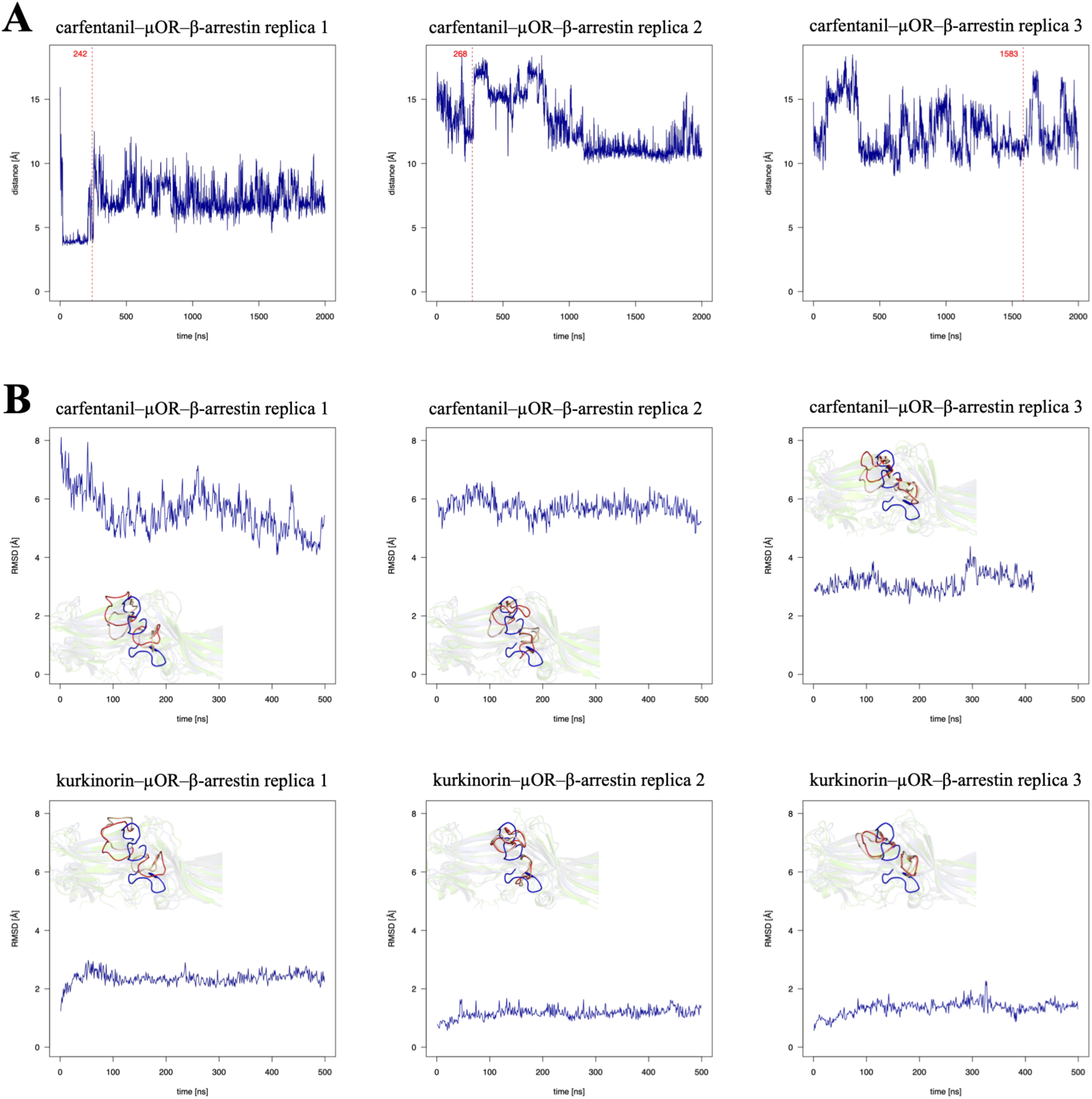
**(A)** The distance between D69^N.s5s6.06^ and R76^N.S6.02^ in the carfentanil–µOR–β-arrestin complexes. The frames exported for use with MGS are indicated in red. **(B)** The heavy-atom RMSD of the finger loop (superposed β-arrestin) for the initial (carfentanil) and ligand-swapped (kurkinorin) µOR–β-arrestin complexes. For the carfentanil replicas, the RMSD was computed starting from the frame exported for MGS. Overlaid are the conformations of the finger and middle loops in the active 7SRS PDB structure (blue), and the first (beige) and last (red) frame of each simulation. Scatter plots of the first 10 PCs are provided in Supplementary Figure S10. The simulation trajectory analysis was limited to the first 500 ns.

The PCA results (**Figure 11**, **Supplementary Figure S4**) again resemble the pattern observed for the µOR–Gi complexes. All three carfentanil-bound replicas show time-ordered, structured PC1/PC2 projections and multimodal histograms, suggesting directed conformational changes that may indicate β-arrestin activation. This is most evident in replica 3, where arcs also appear in higher-order PC pairs, suggesting that rather than a single dominant motion, the protein samples multiple modes. In contrast, the inter-replica variability is greater in the kurkinorin-bound complexes. The more diffuse scatter plots suggest comparatively limited directed conformational change. In kurkinorin replica 3, despite the clear arc shape in the PC1/PC2 projection, its absence in the higher PC pairs suggests a single dominant motion, unlike what was observed in the corresponding carfentanil-bound replica. This confirms the RMSD analysis above and the opposite signaling bias of kurkinorin comparing carfentanil.

**Figure 11.**
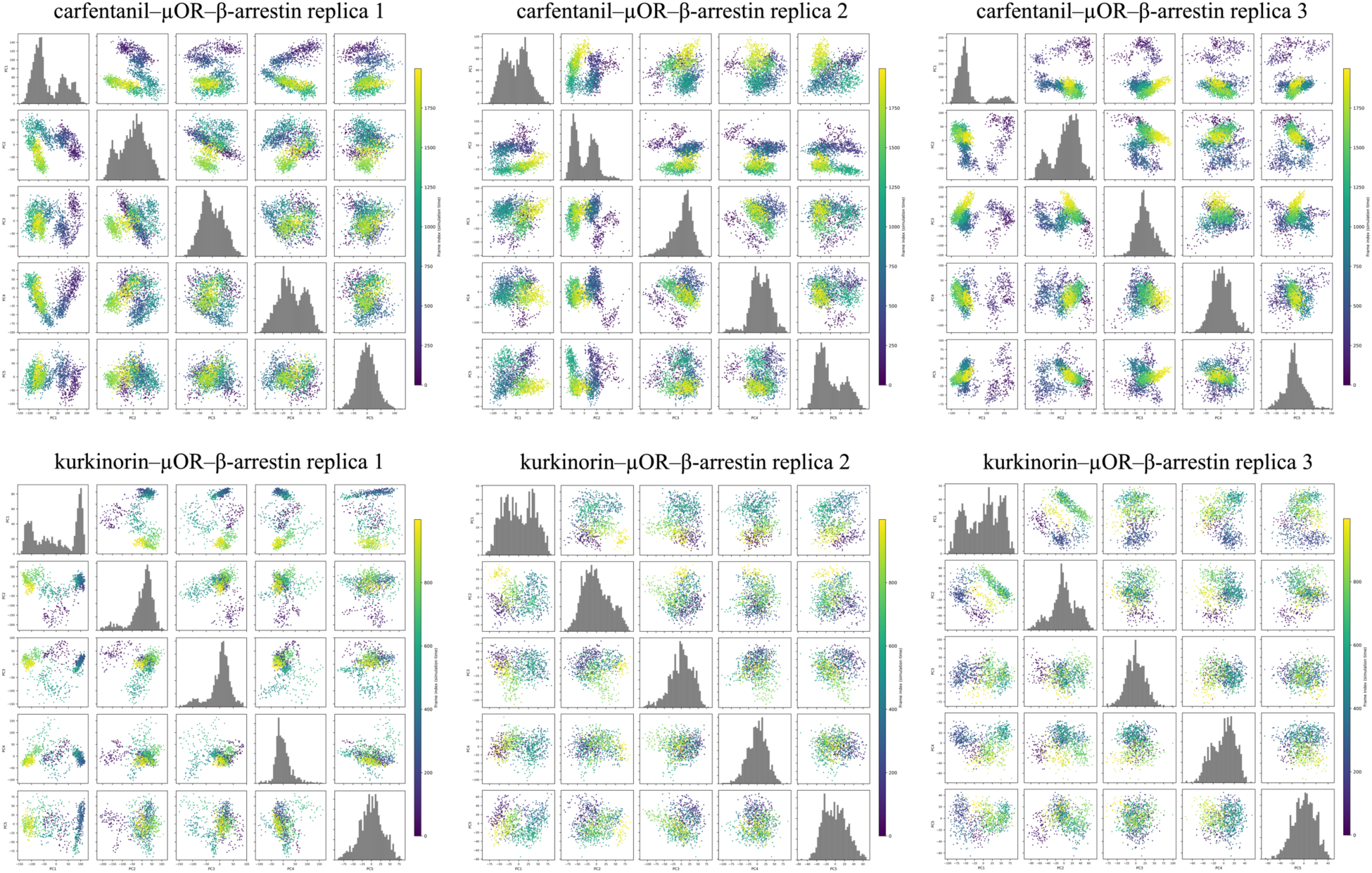
Principal component analysis of the µOR–β-arrestin simulations. Pairwise scatter plots of the projections of the trajectory onto the first 5 PCs, colored by frame index (simulation time; purple = early, yellow = late). Diagonal panels show the distribution of each PC. The analyses were performed on protein backbone atoms following structural alignment.

## 3. Discussion

Here, we showed that 500 ns all-atom, explicit-membrane MD simulations can assess signaling bias of GPCR ligands using microswitch-guided sampling. In at least two of three replicas for each simulated system, clear differences were observed between the initial simulations, where the ligand was biased toward the bound transducer, and ligand-swapped simulations, which involved complexes with oppositely biased ligands. In β_1_AR–Gs complexes, molecular switches II and III were effective for sampling the 2 µs trajectories for MGS but proved less efficient at assessing transducer activation in the 500 ns ligand-test simulations. The heavy-atom RMSD of the G protein and its AHD proved more effective for this task, correctly indicating large-scale conformational changes related to Gs activation within the 500 ns simulation span. In β_1_AR–β-arrestin complexes, activation-like conformational changes were slightly harder to determine, though examining the finger and middle loop conformations yielded an RMSD-based marker for determining β-arrestin activation. Notably, the molecular switches used for MGS sampling of 2 µs trajectories were not necessarily good markers for assessing the activation-like changes of transducers in the 500 ns simulations. It was caused by a substitution of the lipid environment in the ligand-swapped and test simulations, in addition to changes in the ligand. Such lipid substitution may require additional equilibration, typically lasting 10–100 ns (25). Lipid fluctuations are additional factors that alter the simulation system and may modify the conformational landscape (**Figure 12**), in addition to the signaling bias introduced by a newly included ligand. The need to substitute the lipid environment currently stems from technical limitations of the designed pipeline and, in principle, could be overcome to further standardize the computational assay.

**Figure 12.**
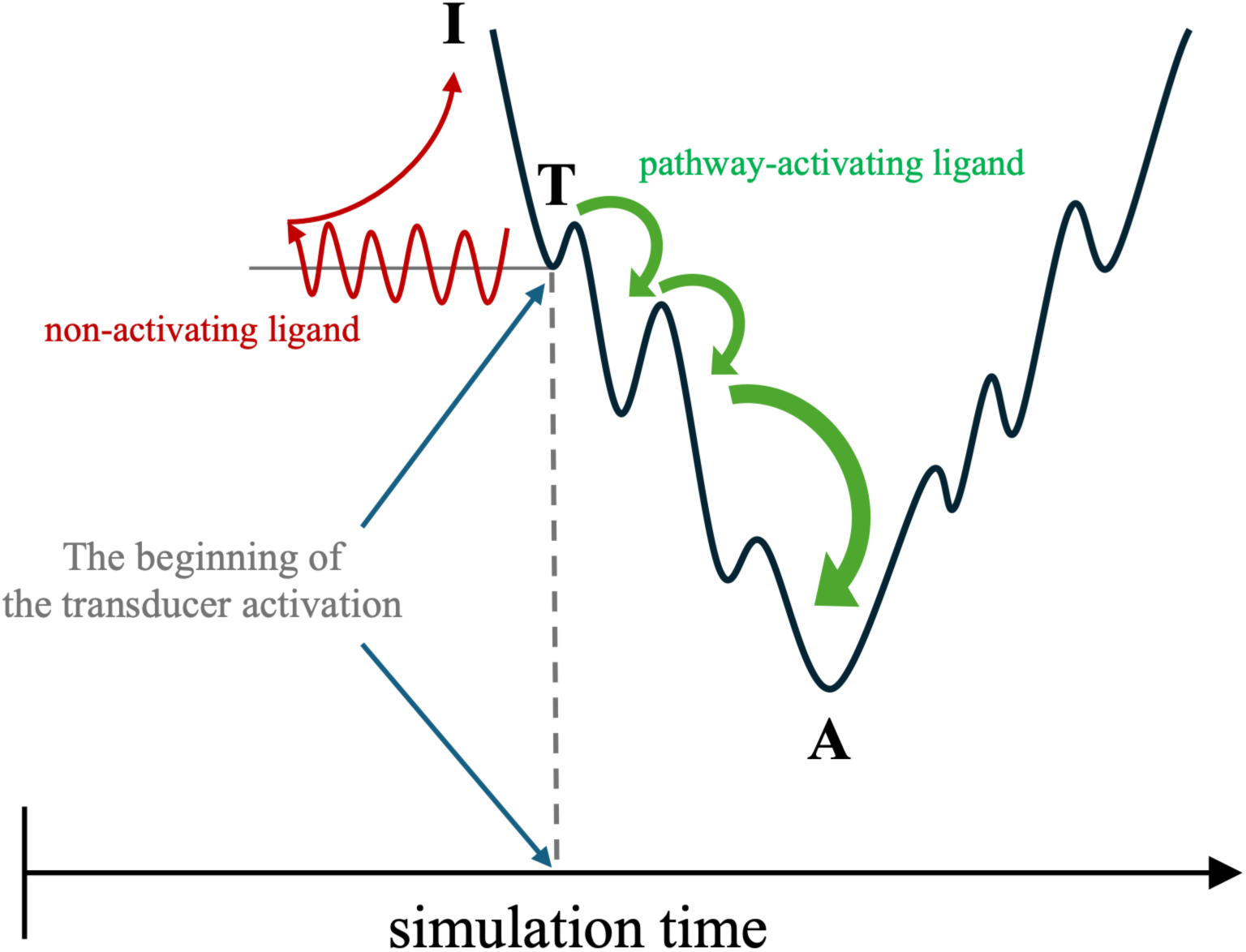
A schematic representation of MGS. A two-dimensional interpretation of the conformational energy landscape of a transducer (I—inactive; T—transition conformation; A—active). MGS frames were exported slightly before (10 ns) the conformational change indicative of transducer activation, as observed in the initial simulation. In the ligand-swapped simulations, a pathway-activating ligand is expected to enable continued progression toward the active state (A), while a non-activating ligand is expected to inhibit further activation, with the complex fluctuating and remaining near the transition conformation (T) or moving toward the inactive conformation.

In summary, these results show that it is possible to predict whether a given β_1_AR ligand activates Gs, β-arrestin, or both. In the CXCR3–Gi systems, the initial and ligand-swapped simulations demonstrated that it is possible to determine whether a ligand activates the Gi signaling pathway. In the µOR–Gs complexes, the results were similarly promising, with the distance between switches II and III identifying frames for MGS and a hydrogen-bond analysis assessing the ongoing G protein activation. As in the β_1_AR–β-arrestin system, activation-like conformational changes within certain microswitches were harder to identify in µOR–β-arrestin complexes than in µOR–Gs complexes, but the RMSD calculations, together with the PCA analysis, unambiguously enabled to determine whether the transducer activation continued. The consistency of the RMSD, hydrogen bond, and PCA results across all biased ligand–GPCR–transducer complexes tested here confirms that MGS could be extended to other GPCR–transducer systems.

Explicit membrane, all-atom MD (AA MD) simulations performed here did not incorporate enhanced sampling methods, e.g., metadynamics. However, we do not consider it as a limitation of a study but as an intended pathway, because for example metadynamics can bias the conformational landscape toward already known transitions (26). It could be counterproductive when the aim is to uncover new conformational changes, especially those involving microswitches. Additionally, metadynamics relies on prior knowledge of the system free energy landscape (26), and no such extensive simulations requiring massive computational resources have been carried out for the complexes described in this study nor for any novel ligands. On the other hand, replica exchange would require more replicas than the three we used in the initial simulations to be effective, further increasing the computational cost (27). Simulated annealing also does not ensure the system finds the global energy minimum and does not prevent it from get trapped in a local minimum (28). Instead, we accelerated our MD simulations using GPUs and, most importantly, sampled trajectories with MGS to find conformations representing local transition-like minima, bypassing the time-consuming process of finding a global minimum with AA MD.

A minor limitation of the current version of MGS is the limited number of well-characterized, fully biased ligands that form the basis of the reference computational assays. For example, for the CXCR3 system, we had to use VUF10661 and VUF11418, which are partially rather than fully biased ligands. Even for extensively studied receptors, only a few fully biased ligands have been identified so far (29). This, in turn, prevents the use of more sophisticated methods, such as machine learning, which could otherwise be used to develop computational assays in a ligand-based approach. To our knowledge, only computational assays for detecting drug target or receptor-type selectivity of tested ligands have been developed using a ligand-based machine-learning approach (30). Nevertheless, the approach presented here does not require large datasets to be effective, but rather a few well-characterized, fully biased ligands for a single receptor, making the ligand signaling bias prediction with computational methods feasible.

These limitations point to several directions for future studies. As more GPCR–transducer structures and larger biased ligand datasets become available, MGS could be extended to additional biased GPCR systems beyond β_1_AR, CXCR3, and µOR. This will allow for further optimization of the method in terms of simulation time, number of replicas, and microswitches used for MGS in both G protein and β-arrestin systems. Although MGS could also incorporate additional enhanced sampling techniques to reduce computational cost, we emphasize the advantage of all-atom simulations of biological systems that resemble the in vivo microenvironment as closely as possible.

## 4. Materials and Methods

### 4.1. Structure Prediction of Ligand–GPCR–Transducer Complexes

Receptors for the biased ligand–GPCR–transducer systems were selected based on the availability of receptors with ligands biased toward either β-arrestin or G proteins, as included in the Biased Signaling Atlas (29) and published research. Another reason for selecting these receptor systems was their direct or indirect connection to immunology and our previous research (31–34). This search yielded three receptors: β_1_AR, CXCR3, and µOR. The receptor systems used in the study included the receptor itself, a biased ligand, and a corresponding transducer (**Table 1**). The receptor complexes were prepared using template-based modeling with MODELLER (35). Template structures were obtained from the PDB after comparing resolution, structural completeness, and the bound transducer in available GPCR structures. To simplify the models, residues C393–D432 of β_1_AR were removed from β_1_AR–β-arrestin complexes, and residues C393–V477 were removed from G protein complexes (UniProt (36) entry P08588). Similarly, residues C338–L369 of CXCR3 (UniProt entry P49682) and residues T356–P400 of µOR (UniProt entry P35372) were removed from their respective G protein complexes. Missing loops were reconstructed using the MODELLER DOPEHRLoopModel method.

**Table 1.**
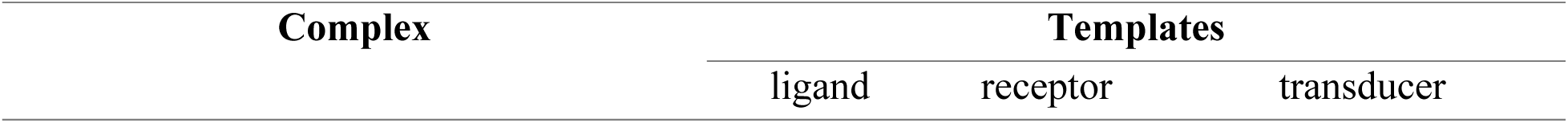
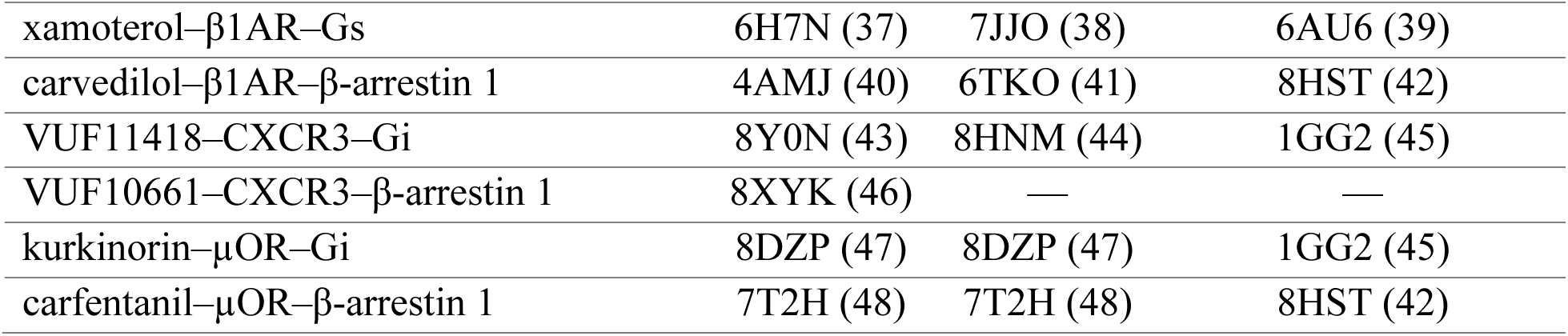
The PDB entries used as templates for the receptor complex structure prediction.

Some ligands, such as carfentanil and kurkinorin, were absent from all available PDB receptor structures. Accordingly, structures containing very similar ligands and assumed similar binding modes were used, and the ligands were modified into the intended compounds in Maestro. This was followed by gradient-based LBFGS minimization using the default OPLS force field to remove steric clashes with side-chain rotamers (49). Molecular docking was not used here, as it would introduce additional uncertainty in pose selection. For each system, 500 initial receptor models with ligands and transducers, along with 10 loop models, were generated with MODELLER, yielding 5000 models per receptor complex. 200 lowest-energy models by DOPE scores were evaluated in PyMOL (50), and for each complex, three models with the lowest RMSD relative to the templates were selected for three MD simulation replicas. Hydrogen atoms were added to the models using the Protein Preparation Workflow in Maestro (49).

### 4.2. Molecular Dynamics Simulations

Inputs for the MD simulations were prepared using the aforementioned models of complexes and CHARMM-GUI’s (51, 52) Membrane Builder (53). Disulfide bridges were defined based on the PDB templates, and the following residues were phosphorylated in accordance with current knowledge of these receptor systems: S459, S461, and S462 for β_1_AR (54); T372, S377, T378, and T381 for µOR (55). Ligand parameters were determined using the CHARMM General Force Field (CGenFF). The receptor complexes were inserted into a lipid bilayer composed of 1-palmitoyl-2-oleoyl-*sn*-glycero-3-phosphocholine (POPC) and cholesterol at a 3:1 ratio (13, 14). K^+^ and Cl^−^ ions were added at a concentration of 0.15 M and a rectangular TIP3 water box was generated. After assembling the systems, additional CHARMM minimization (56) was performed before generating MD input files.

MD simulations were performed using NAMD 3.0 with GPU acceleration (57, 58) and the CHARMM36m force field (59). Equilibration began with 10000 steps of steepest descent minimization followed by 25000 steps of conjugate gradient minimization. The system was equilibrated in the NVT ensemble using Langevin dynamics at 303.15 K. A 2 fs timestep was used for both the equilibration and production runs. Production runs were carried out in the NPT ensemble using the Langevin piston Nosé–Hoover method at 1 bar and 303.15 K and lasted between 2 µs and 2.5 µs. Each system was simulated in three replicas. Every tenth trajectory frame was combined using CatDCD (60) and wrapped using PBCTools (61). Analyses were performed using a combination of VMD (62), PyMOL, and the MDAnalysis Python library (63), focusing on conformational changes involving microswitches.

### 4.3. Microswitch-Guided Sampling

Molecular switches (micro- and macroswitches, if differentiated as in (64)) identified in the initial 2–2.5 µs simulations were used to determine activation-like transition points for trajectory sampling and extract representative complex conformations (**Table 2**). These transition-like conformations were used as the basis for the receptor-specific computational assay developed to assess the functional selectivity of GPCR ligands. Although some microswitches were similar across receptor complexes, many differed due to the use of different G protein subtypes, e.g., Gs in β_1_AR and Gi in CXCR3 and µOR complexes. Notably, none of the examined microswitches were in receptors, as these were expected to remain in an active conformation throughout the simulations regardless of which signaling pathway a tested agonist was biased to. However, active conformations of receptors binding differently biased ligands should also differ, and so should receptor microswitches. Here, only molecular switches in the transducers were examined to observe activation-like or deactivation-like changes. Differences between transducers and the resulting variation in microswitch choice led to differences in the time points guiding trajectory sampling. In addition, MD simulation replicas starting from different complex conformations differ in the timing of a microswitch change, because MD simulations tend to become trapped in local minima (65). Nevertheless, these differences in microswitches and the time points chosen for trajectory sampling could be considered an optimization of the receptor- and transducer-specific computational assay for a specific GPCR system. In other words, the computational assay developed for β_1_AR in this study cannot be used to test ligands of other GPCRs. The limitations of molecular dynamics, such as trapping simulations in local minima, are not relevant to the developed computational assays because they do not aim to depict a fully activated signaling pathway as experimental assays do. GPCR signal transduction occurs through a series of conformational changes involving the receptor (12, 66) and transducers (13, 34, 67, 68). Our MGS method builds on this foundation by focusing on local transition states that contribute to the activation of the ligand-receptor-transducer complex (**Figure 12**).

**Table 2.** MGS implementation in the performed MD simulations.

| Reference ligand–<br>receptor complex | Transducer | Microswitch* | Activation-like change** | Extracted frame [ns] |  |  |
| --- | --- | --- | --- | --- | --- | --- |
|  |  |  |  | replica 1 | replica 2 | replica 3 |
| xamoterol–$\beta_1$AR | Gs | II and III | Increase in the distance between Cζ of R228<sup>G.H2.01</sup> and Cδ of E259<sup>G.4h3.10</sup> | 1637 | 1867 | 1472 |
| carvedilol–$\beta_1$AR | β-arrestin 1 | finger loop | Increase in the finger loop RMSD | 872 | 181 | 762 |
| VUF11418–CXCR3 | Gi | D295<sup>G.HG.02</sup> | Changes in the $\chi_1$ dihedral angle | 498 | 1001 | 1162 |
| kurkinorin–μOR | Gi | II and III | Increase in the distance between Cζ of R205<sup>G.H2.01</sup> and Cδ of E236<sup>G.4h3.10</sup> | 294 | 622 | 1162 |
| carfentanil–μOR | β-arrestin 1 | finger loop | Increase in the distance between D69<sup>N.s5s6.06</sup> and R<sup>76N.S6.02</sup> | 242 | 268 | 1583 |

\*Microswitch conformational changes are shown in **Figure 2A**, **Figure 4A**, **Figure 6A**, **Figure 8A**, and **Figure 10A**.
\*\*The opposite changes were observed for ligands biased toward the other transducer.

For each replica of the initial simulations, a representative, transition-like conformation of the complex was extracted from the 10^th^ frame before the microswitch-guided time point. This 10 ns interval, as close as possible to the activation-like change of the microswitch, aimed to account for lipid relaxation (equilibration) in newly ligand-swapped complex systems (25), while providing ample time for the microswitch to undergo a conformational change.

### 4.4. Ligand-Swapped Complexes

A second set of simulations involved complexes in which the ligand was biased toward the opposite transducer (ligand-swapped complexes). Rather than starting simulations anew, a single frame was exported from each original trajectory, sampled 10 ns before an observed active-like transition of at least one identified microswitch. Then, ligand structure modifications in Maestro were used to replace the original ligand with a ligand biased toward the opposite signaling pathway, followed by gradient-based LBFGS minimization with the default OPLS force field to remove steric clashes with side-chain rotamers (49). Simulation inputs for NAMD 3.0 were prepared as described above using CHARMM-GUI’s (51, 52) Membrane Builder (53), with the simulations lasting 1 µs. However, to meet the short computational time required to test multiple ligands in a computational assay, we analyzed only the first 500 ns. This series of ligand-swapped simulations aimed to determine whether replacing the biased ligand with the oppositely biased ligand could prevent the microswitches from changing to an active-like conformation.

### 4.5. Complexes of Tested Ligands

We performed a third set of simulations for β_1_AR, replacing the ligands with other known biased ligands—nebivolol and isoetharine—following the procedure described for the ligand-swapped complexes. The simulations lasted 500 ns for reasons mentioned above and aimed to test the signaling bias of these ligands. We again assessed whether microswitch conformational changes were similar to those observed in simulations with the reference biased ligands. Similar conformational changes suggested similar signaling bias in the tested ligands, while no conformational changes or deactivation-like changes suggested the opposite signaling bias. In the G protein complexes, signaling bias was assessed by examining RMSD values of G*a* and its α-helical domain, as well as the formation of hydrogen bonds between R205^G.H2.01^ and E236^G.4h3.10^. In the β-arrestin systems, RMSD of the finger loop, hydrogen bonds, and loop conformations relevant to β-arrestin activation were examined instead.

## Acknowledgements

The first version of the microswitch-guided sampling (MGS) method was submitted as an application for PRELUDIUM 24 (ID 644644), National Science Centre (NCN) in Poland, in June 2025. We gratefully acknowledge the Polish high-performance computing infrastructure PLGrid (HPC Centers: ACK Cyfronet AGH) for providing computer facilities and support within computational grant no. PLG/2026/019390. This work received no external funding. No author has an actual or perceived conflict of interest with the contents of this article.

## Data Availability Statement

All data generated or analyzed during this study are included in this published article, its Supplementary Information file S1.pdf and deposited in Zenodo.

## Author contributions

Conceptualization: PD, DL; data curation: PD, DL; formal analysis: PD, DL; funding acquisition: DL; investigation: PD, DL; methodology: PD, DL; project administration: PD, DL; resources: DL; supervision: DL; validation: PD, DL; visualization: PD, DL; writing – original draft: PD, DL; writing – review and editing: PD, DL.

## References

1. P. Kolb, et al., Community guidelines for GPCR ligand bias: IUPHAR review 32. British Journal of Pharmacology 179, 3651–3674 (2022).

2. R. Alhadeff, I. Vorobyov, H. W. Yoon, A. Warshel, Exploring the free-energy landscape of GPCR activation. PNAS 115, 10327–10332 (2018).

3. Z. Xu, Z. Shao, Dynamic mechanism of GPCR-mediated β-arrestin: a potential therapeutic agent discovery of biased drug. Sig Transduct Target Ther 7, 283 (2022).

4. M. Bermudez, T. N. Nguyen, C. Omieczynski, G. Wolber, Strategies for the discovery of biased GPCR ligands. Drug Discovery Today 24, 1031–1037 (2019).

5. M. Ippolito, J. L. Benovic, Biased agonism at β-adrenergic receptors. Cellular Signalling 80, 109905 (2021).

6. P. Dragan, D. Latek, The two-sided impact of beta-adrenergic receptor ligands on inflammation. Current Opinion in Physiology 41, 100779 (2024).

7. F. J. Ehlert, H. Suga, M. T. Griffin, Analysis of Agonism and Inverse Agonism in Functional Assays with Constitutive Activity: Estimation of Orthosteric Ligand Affinity Constants for Active and Inactive Receptor States. The Journal of Pharmacology and Experimental Therapeutics 338, 671–686 (2011).

8. D. Wootten, A. Christopoulos, M. Marti-Solano, M. M. Babu, A. Et., Mechanisms of signalling and biased agonism in G protein-coupled receptors. Nature Reviews Molecular Cell Biology (2018). 10.1038/s41580-018-0049-3.

9. Y. Ma, B. Patterson, L. Zhu, Biased signaling in GPCRs: Structural insights and implications for drug development. Pharmacology & Therapeutics 266, 108786 (2025).

10. M. Lopez-Balastegui, et al., Relevance of G protein-coupled receptor (GPCR) dynamics for receptor activation, signalling bias and allosteric modulation. British Journal of Pharmacology 182, 3211–3224 (2025).

11. B. Trzaskowski, et al., Action of Molecular Switches in GPCRs - Theoretical and Experimental Studies. CMC 19, 1090–1109 (2012).

12. A. S. Hauser, et al., GPCR activation mechanisms across classes and macro/microscales. Nat Struct Mol Biol 28, 879–888 (2021).

13. P. Dragan, V. Gratio, T. Voisin, A. Couvineau, D. Latek, Non-canonical Gq activation by orexin receptor type 2 and lemborexant observed in microsecond molecular dynamics simulations. Sci Rep 15, 30899 (2025).

14. V. Gratio, et al., Pharmacodynamics of the orexin type 1 (OX1) receptor in colon cancer cell models: A two-sided nature of antagonistic ligands resulting from partial dissociation of Gq. British Journal of Pharmacology 182, 1528–1545 (2025).

15. O. Fleetwood, P. Matricon, J. Carlsson, L. Delemotte, Energy Landscapes Reveal Agonist Control of G Protein-Coupled Receptor Activation via Microswitches. Biochemistry 59, 880–891 (2020).

16. S. Della Longa, A. Arcovito, Microswitches for the Activation of the Nociceptin Receptor Induced by Cebranopadol: Hints from Microsecond Molecular Dynamics. J. Chem. Inf. Model. 59, 818–831 (2019).

17. T. Che, H. Dwivedi-Agnihotri, A. K. Shukla, B. L. Roth, Biased ligands at opioid receptors: Current status and future directions. Science Signaling 14, eaav0320 (2021).

18. M. Ippolito, J. L. Benovic, Biased agonism at β-adrenergic receptors. Cell Signal 80, 109905 (2021).

19. V. Lukasheva, et al., Signal profiling of the β1AR reveals coupling to novel signalling pathways and distinct phenotypic responses mediated by β1AR and β2AR. Sci Rep 10, 8779 (2020).

20. K. Zheng, et al., Biased agonists of the chemokine receptor CXCR3 differentially signal through Gαi:β-arrestin complexes. Science Signaling 15, eabg5203 (2022).

21. A. Faouzi, B. R. Varga, S. Majumdar, Biased Opioid Ligands. Molecules 25, 4257 (2020).

22. N. Ramos-Gonzalez, et al., Carfentanil is a β-arrestin-biased agonist at the μ opioid receptor. British Journal of Pharmacology 180, 2341–2360 (2023).

23. Sente, Peer, Srivastava, Baidya, A. Et., Molecular mechanism of modulating arrestin conformation by GPCR phosphorylation. Nature Structural & Molecular Biology (2018). 10.1038/s41594-018-0071-3.

24. H. Zhang, et al., The molecular basis of μ-opioid receptor signaling plasticity. Cell Res 35, 1021–1036 (2025).

25. D. J. Smith, J. B. Klauda, A. J. Sodt, Simulation Best Practices for Lipid Membranes [Article v1.0]. Living J Comput Mol Sci 1, 5966 (2019).

26. G. Bussi, A. Laio, Using metadynamics to explore complex free-energy landscapes. Nat Rev Phys 2, 200–212 (2020).

27. X. Gong, Y. Zhang, J. Chen, Advanced Sampling Methods for Multiscale Simulation of Disordered Proteins and Dynamic Interactions. Biomolecules 11, 1416 (2021).

28. S. Kannan, M. Zacharias, Simulated annealing coupled replica exchange molecular dynamics—An efficient conformational sampling method. Journal of Structural Biology 166, 288–294 (2009).

29. J. Caroli, et al., A community Biased Signaling Atlas. Nat Chem Biol 19, 531–535 (2023).

30. D. Latek, K. Prajapati, P. Dragan, M. Merski, P. Osial, GPCRVS - AI-driven Decision Support System for GPCR Virtual Screening. International Journal of Molecular Sciences 26, 2160 (2025).

31. S. G. Sanmukh, et al., “Is Cancer Our Equal or Our Better? Artificial Intelligence in Cancer Drug Discovery” in Artificial Intelligence and Bioinformatics in Cancer: An Interdisciplinary Approach, N. Rezaei, Ed. (Springer Nature Switzerland, 2025), pp. 223–258.

32. P. Dragan, M. Merski, S. Wiśniewski, S. G. Sanmukh, D. Latek, Chemokine Receptors— Structure-Based Virtual Screening Assisted by Machine Learning. Pharmaceutics 15, 516 (2023).

33. P. Dragan, K. Joshi, A. Atzei, D. Latek, Keras/TensorFlow in Drug Design for Immunity Disorders. IJMS 24, 15009 (2023).

34. P. Dragan, D. Latek, C5aR2 signaling mechanisms via the β-arrestin-dependent pathway observed in microsecond-scale MD simulations. The Journal of Pharmacology and Experimental Therapeutics 105028 (2026). 10.1016/j.jpet.2026.105028.

35. N. Eswar, et al., Comparative Protein Structure Modeling Using Modeller. CP in Bioinformatics 15 (2006).

36. The UniProt Consortium, UniProt: the Universal Protein Knowledgebase in 2025. Nucleic Acids Research 53, D609–D617 (2025).

37. T. Warne, P. C. Edwards, A. S. Doré, A. G. W. Leslie, C. G. Tate, Molecular basis for high affinity agonist binding in GPCRs. [Preprint] (2018). Available at: http://biorxiv.org/lookup/doi/10.1101/436212 [Accessed 27 August 2026].

38. M. Su, et al., Structural Basis of the Activation of Heterotrimeric Gs-Protein by Isoproterenol-Bound β1-Adrenergic Receptor. Molecular Cell 80, 59–71.e4 (2020).

39. Q. Hu, K. M. Shokat, Disease-Causing Mutations in the G Protein Gαs Subvert the Roles of GDP and GTP. Cell 173, 1254–1264.e11 (2018).

40. T. Warne, P. C. Edwards, A. G. W. Leslie, C. G. Tate, Crystal Structures of a Stabilized β1-Adrenoceptor Bound to the Biased Agonists Bucindolol and Carvedilol. Structure 20, 841–849 (2012).

41. Y. Lee, et al., Molecular basis of β-arrestin coupling to formoterol-bound β1-adrenoceptor. Nature 583, 862–866 (2020).

42. Y. Yun, et al., GPCR targeting of E3 ubiquitin ligase MDM2 by inactive β-arrestin. Proceedings of the National Academy of Sciences 120, e2301934120 (2023).

43. S. Saha, et al., Structural visualization of small molecule recognition by CXCR3 uncovers dual-agonism in the CXCR3-CXCR7 system. Nat Commun 16, 3047 (2025).

44. H. Jiao, et al., Structural insights into the activation and inhibition of CXC chemokine receptor 3. Nat Struct Mol Biol 31, 610–620 (2024).

45. M. A. Wall, et al., The structure of the G protein heterotrimer Giα1β1γ2. Cell 83, 1047–1058 (1995).

46. S. Saha, et al., Structural visualization of small molecule recognition by CXCR3 uncovers dual-agonism in the CXCR3-CXCR7 system. Nat Commun 16, 3047 (2025).

47. J. Han, et al., Ligand and G-protein selectivity in the κ-opioid receptor. Nature 617, 417–425 (2023).

48. Q. Qu, et al., Insights into distinct signaling profiles of the µOR activated by diverse agonists. Nat Chem Biol 19, 423–430 (2023).

49. Schrödinger Release 2021-4: Maestro.

50. L. Schrödinger, W. DeLano, PyMOL. (2020). Deposited 20 May 2020.

51. S. Jo, T. Kim, V. G. Iyer, W. Im, CHARMM-GUI: A web-based graphical user interface for CHARMM. Journal of Computational Chemistry 29, 1859–1865 (2008).

52. B. R. Brooks, et al., CHARMM: The biomolecular simulation program. Journal of Computational Chemistry 30, 1545–1614 (2009).

53. E. L. Wu, et al., CHARMM-GUI Membrane Builder toward realistic biological membrane simulations. Journal of Computational Chemistry 35, 1997–2004 (2014).

54. L. Hinz, A. Ahles, B. Ruprecht, B. Küster, S. Engelhardt, Two serines in the distal C-terminus of the human ß1-adrenoceptor determine ß-arrestin2 recruitment. PLoS ONE 12, e0176450 (2017).

55. O. Underwood, et al., Key phosphorylation sites for robust β-arrestin2 binding at the MOR revisited. Commun Biol 7, 933 (2024).

56. K. Vanommeslaeghe, et al., CHARMM general force field: A force field for drug-like molecules compatible with the CHARMM all-atom additive biological force fields. Journal of Computational Chemistry 31, 671–690 (2010).

57. J. C. Phillips, et al., Scalable molecular dynamics with NAMD. J Comput Chem 26, 1781–1802 (2005).

58. J. C. Phillips, et al., Scalable molecular dynamics on CPU and GPU architectures with NAMD. J. Chem. Phys. 153, 044130 (2020).

59. J. Huang, et al., CHARMM36m: an improved force field for folded and intrinsically disordered proteins. Nat Methods 14, 71–73 (2017).

60. J. Gullingsrud, CatDCD. (version 4.0). Deposited version 4.0.

61. T. Giorgino, J. Henin, O. Lenz, C. Mura, J. Saam, PBCTools. (version 2.7). Deposited version 2.7.

62. W. Humphrey, A. Dalke, K. Schulten, VMD: Visual molecular dynamics. Journal of Molecular Graphics 14, 33–38 (1996).

63. R. J. Gowers, et al., MDAnalysis: A Python Package for the Rapid Analysis of Molecular Dynamics Simulations. SciPy 2016 (2016). 10.25080/Majora-629e541a-00e.

64. A. S. Hauser, et al., GPCR activation mechanisms across classes and macro/microscales. Nature Structural & Molecular Biology 28, 879–888 (2021).

65. D. Van Der Spoel, M. M. Seibert, Protein Folding Kinetics and Thermodynamics from Atomistic Simulations. Phys. Rev. Lett. 96, 238102 (2006).

66. B. K. Kobilka, X. Deupi, Conformational complexity of G-protein-coupled receptors. Trends in Pharmacological Sciences 28, 397–406 (2007).

67. D. Ham, et al., Conformational switch that induces GDP release from Gi. J Struct Biol 213, 107694 (2021).

68. J. S. Smith, S. Rajagopal, The β-Arrestins: Multifunctional Regulators of G Protein-coupled Receptors. J Biol Chem 291, 8969–8977 (2016).

