## Supplementary information file S1.pdf for "Microswitch-Guided Sampling for the Detection of Ligand Signaling Bias in GPCR Systems"

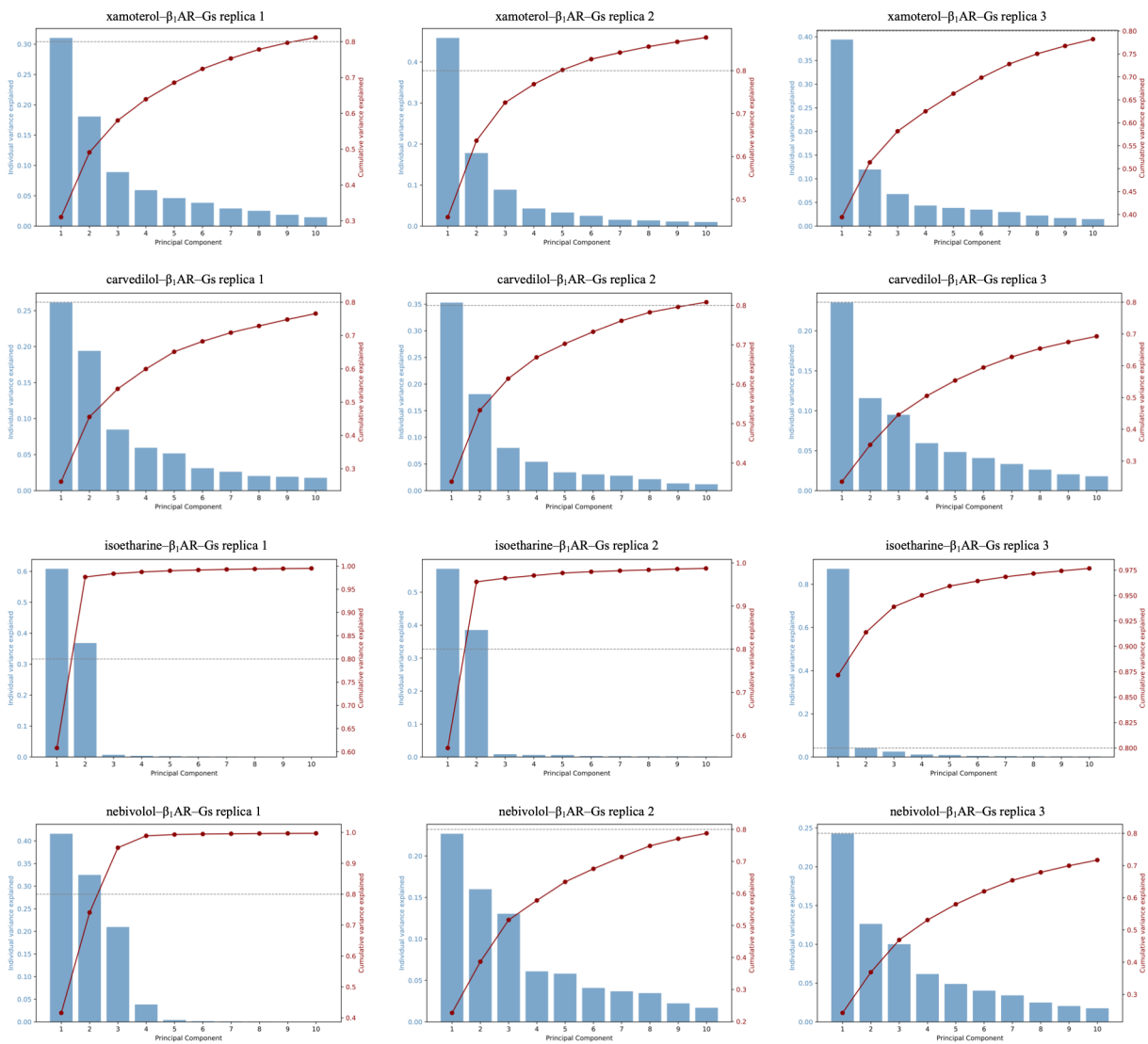

**Figure S1. Scree plots of principal component analysis of the $\beta_1$AR-Gs simulations.** Bars (left axis, blue) show the fraction of total variance explained by each of the first 10 principal components. The analyses were performed on protein backbone atoms following structural alignment.

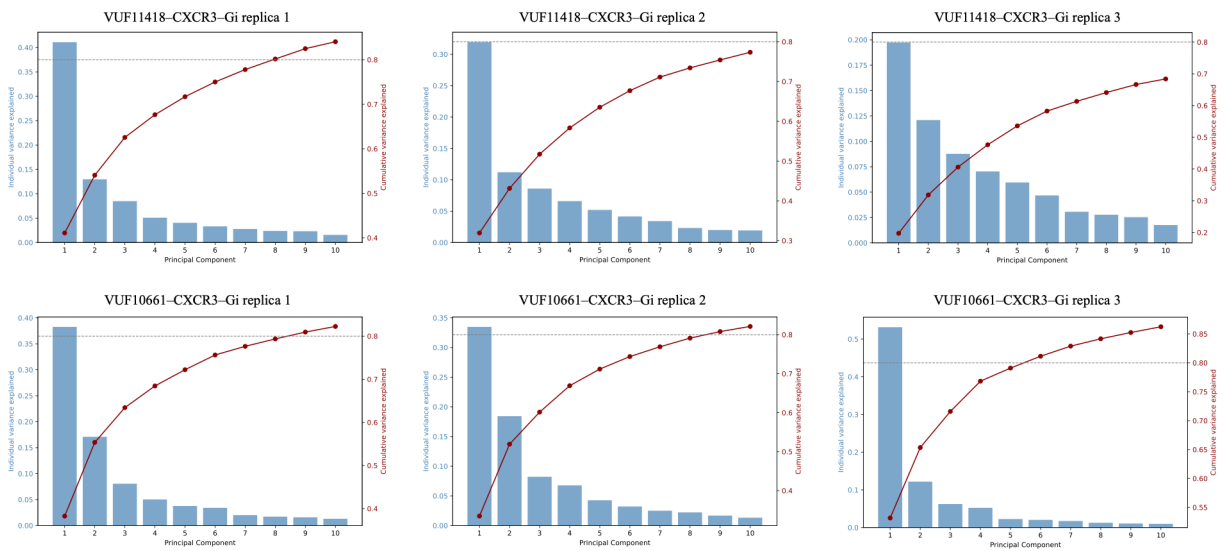

**Figure S2. Scree plots of principal component analysis of the CXCR3-Gi simulations.** Bars (left axis, blue) show the fraction of total variance explained by each of the first 10 principal components. The analyses were performed on protein backbone atoms following structural alignment.

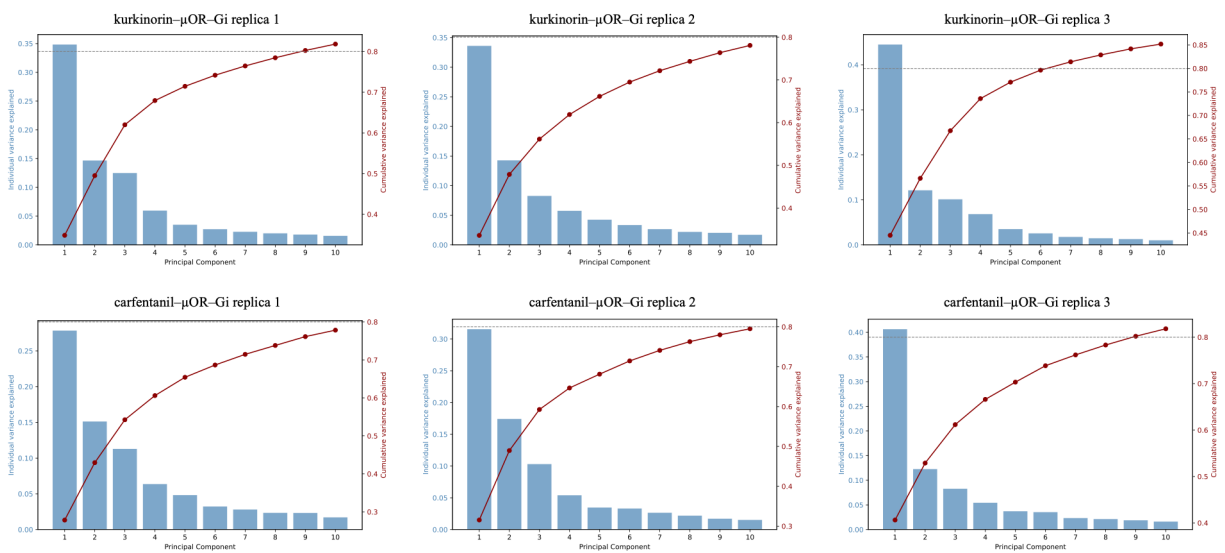

**Figure S3. Scree plots of principal component analysis of the μOR-Gi simulations.** Bars (left axis, blue) show the fraction of total variance explained by each of the first 10 principal components. The analyses were performed on protein backbone atoms following structural alignment.

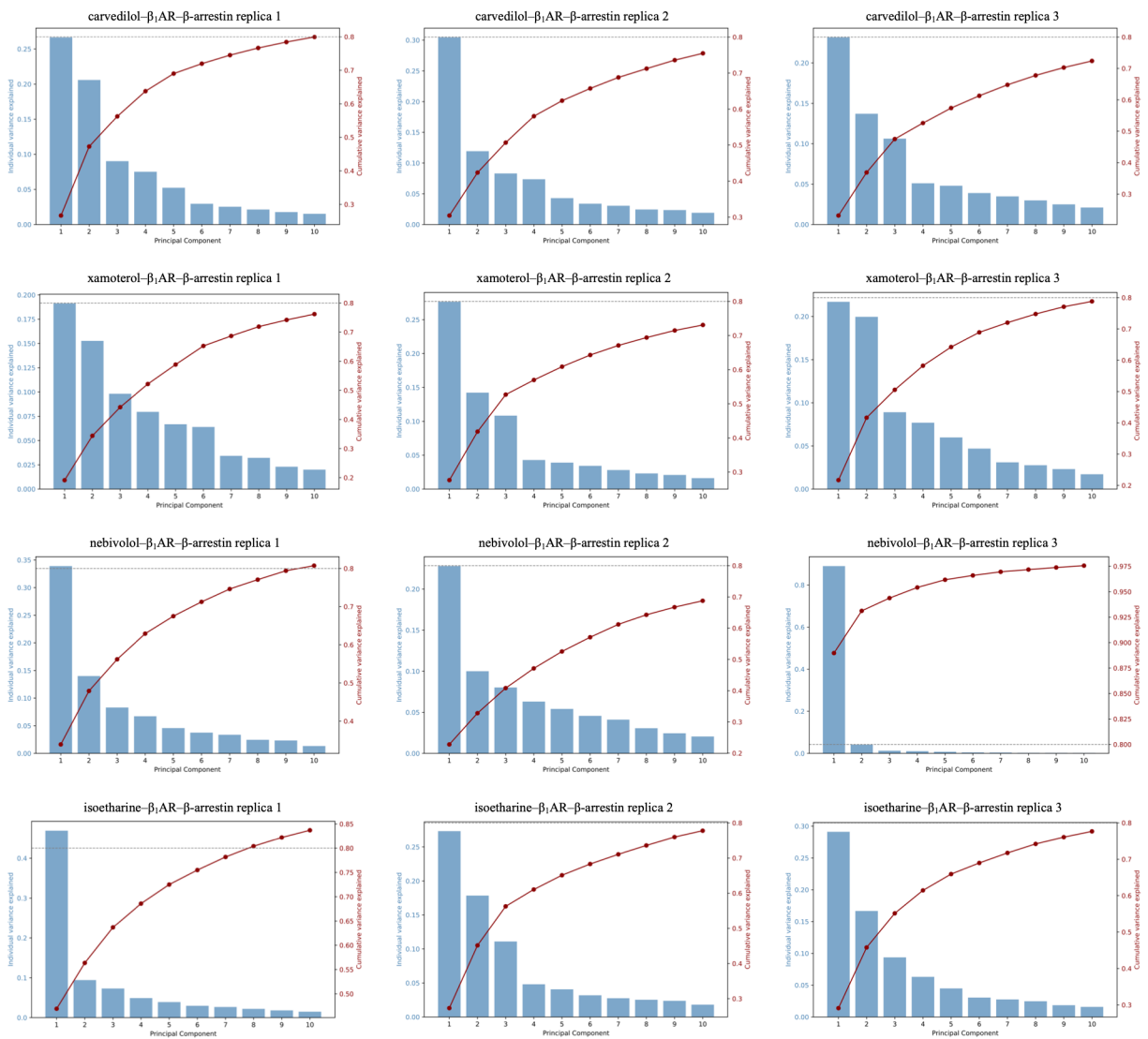

**Figure S4. Scree plots of principal component analysis of the $\beta_1\text{AR}$–β-arrestin simulations.** Bars (left axis, blue) show the fraction of total variance explained by each of the first 10 principal components. The analyses were performed on protein backbone atoms following structural alignment.

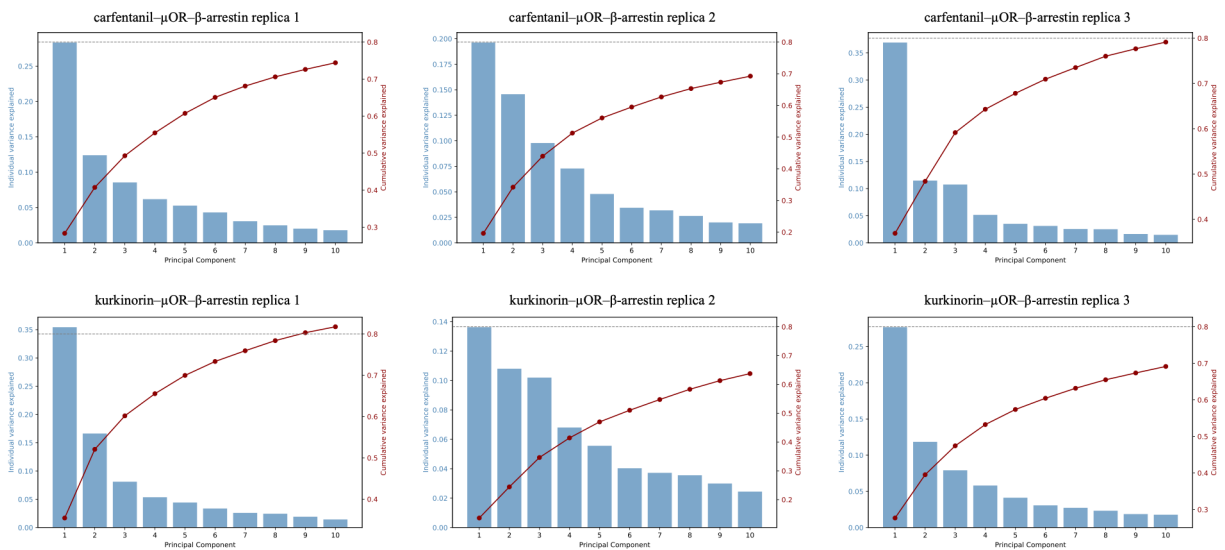

**Figure S5. Scree plots of principal component analysis of the μOR-β-arrestin simulations.** Bars (left axis, blue) show the fraction of total variance explained by each of the first 10 principal components. The analyses were performed on protein backbone atoms following structural alignment

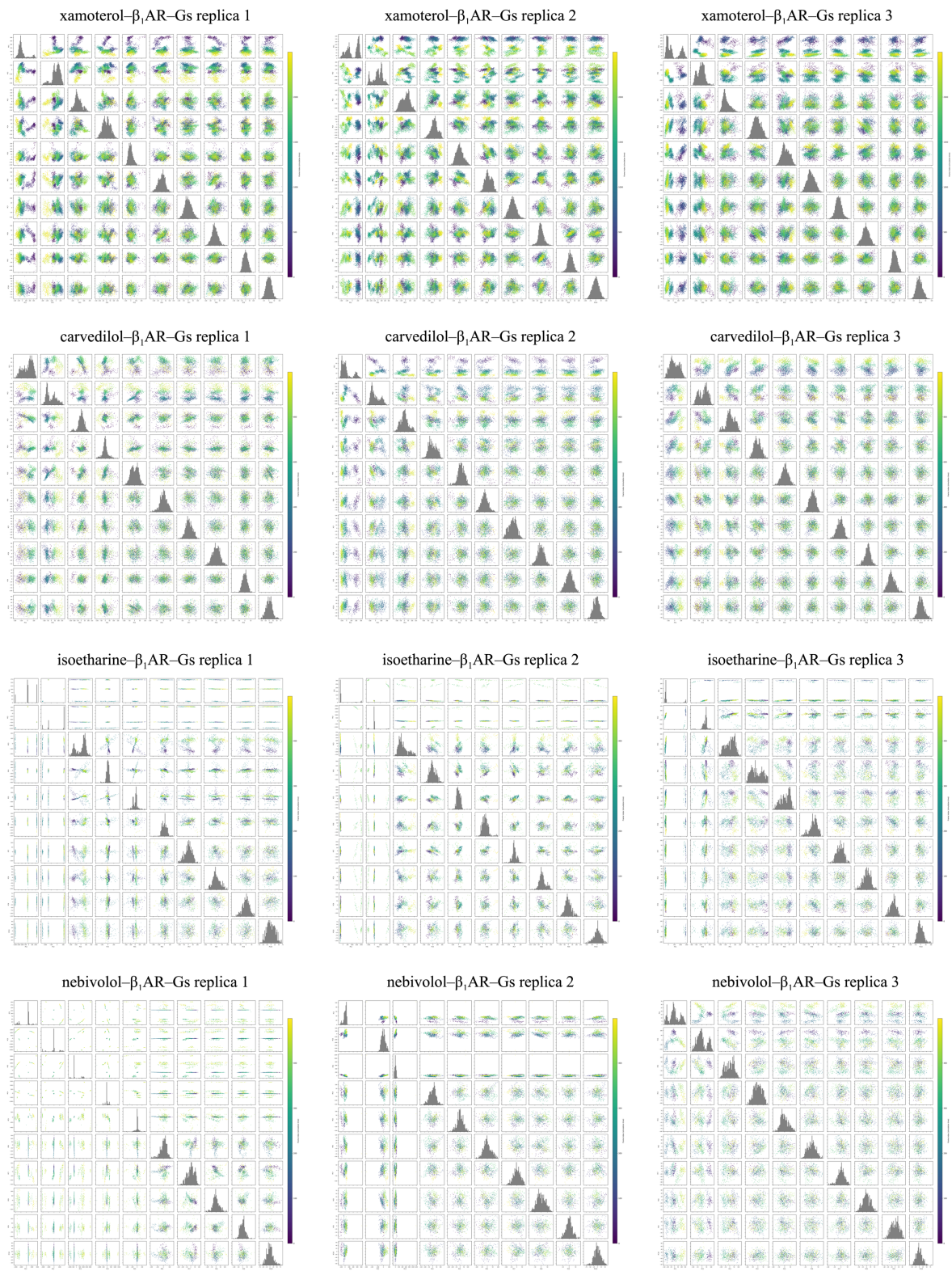

**Figure S6. Principal component analysis of the $\beta_1$AR-Gs simulations.** Pairwise scatter plots of the projections of the trajectory onto the first 10 PCs, colored by frame index (simulation time; purple = early, yellow = late). Diagonal panels show the distribution of each PC. The analyses were performed on protein backbone atoms following structural alignment.

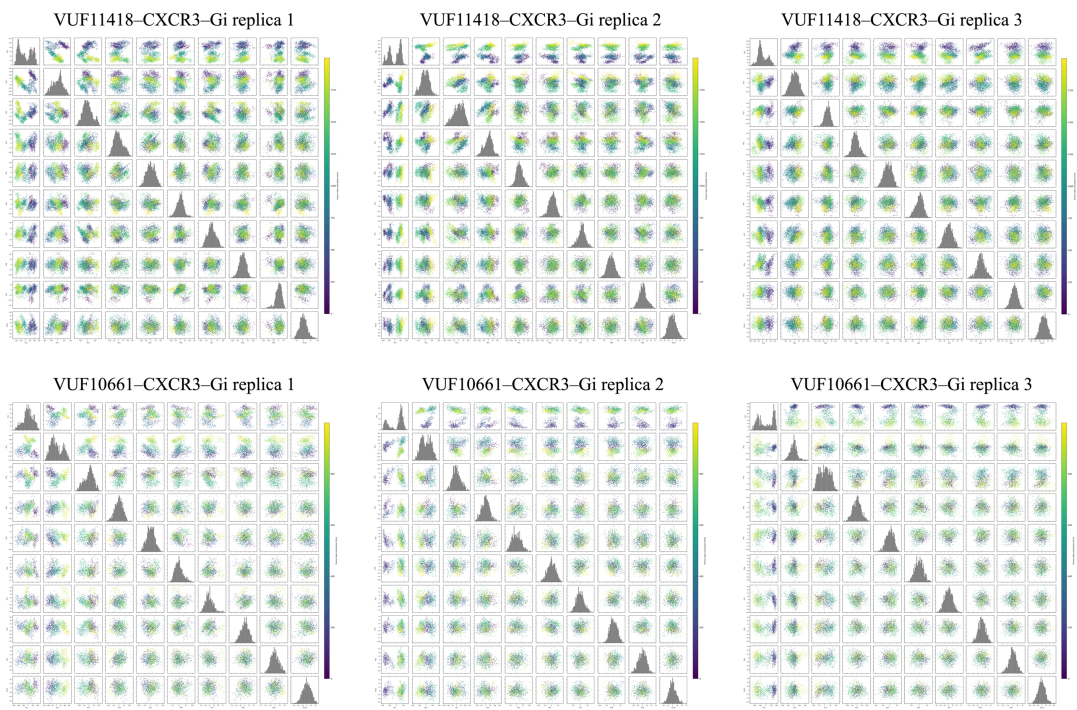

**Figure S7. Principal component analysis of the CXCR3-Gi simulations.** Pairwise scatter plots of the projections of the trajectory onto the first 10 PCs, colored by frame index (simulation time; purple = early, yellow = late). Diagonal panels show the distribution of each PC. The analyses were performed on protein backbone atoms following structural alignment.

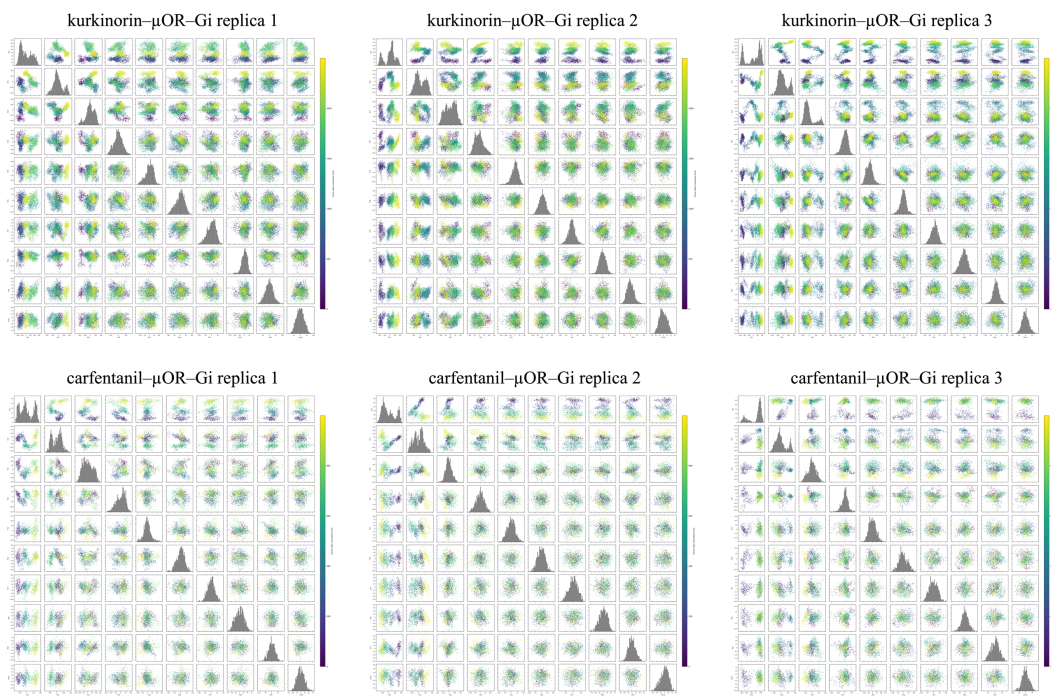

**Figure S8. Principal component analysis of the μOR-Gi simulations.** Pairwise scatter plots of the projections of the trajectory onto the first 10 PCs, colored by frame index (simulation time; purple = early, yellow = late). Diagonal panels show the distribution of each PC. The analyses were performed on protein backbone atoms following structural alignment.

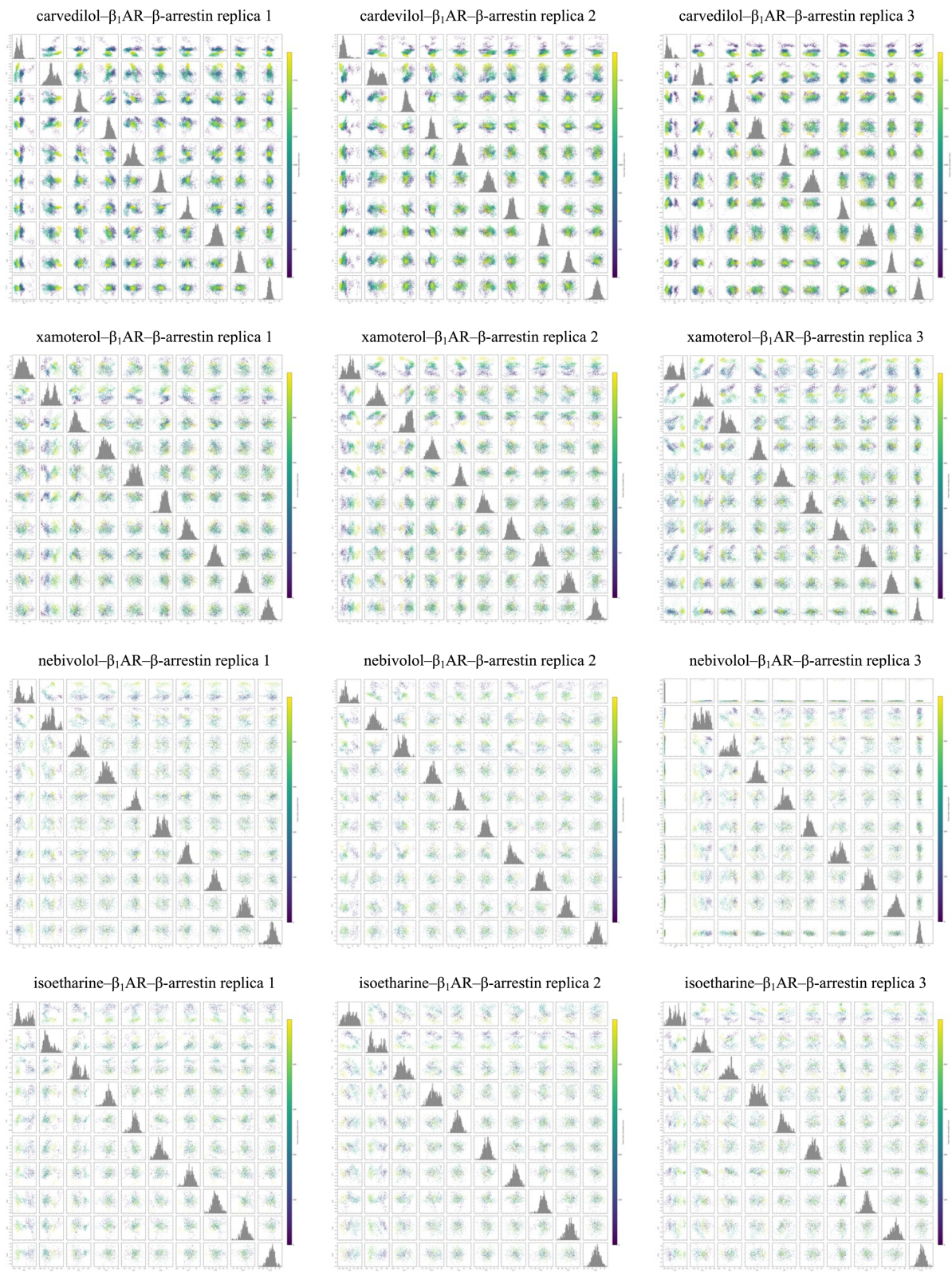

**Figure S9. Principal component analysis of the $\beta_1$AR-β-arrestin simulations.** Pairwise scatter plots of the projections of the trajectory onto the first 10 PCs, colored by frame index (simulation time; purple = early, yellow = late). Diagonal panels show the distribution of each PC. The analyses were performed on protein backbone atoms following structural alignment.

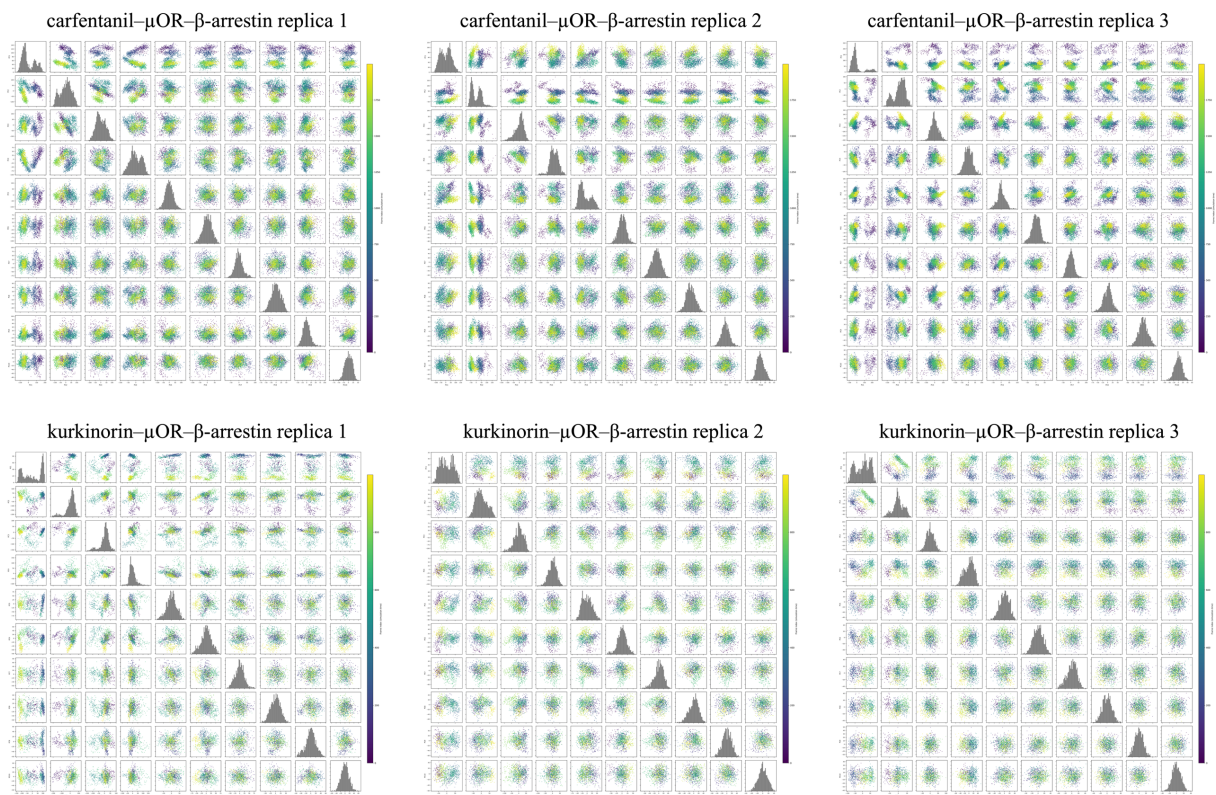

**Figure S10. Principal component analysis of the μOR-β-arrestin simulations.** Pairwise scatter plots of the projections of the trajectory onto the first 10 PCs, colored by frame index (simulation time; purple = early, yellow = late). Diagonal panels show the distribution of each PC. The analyses were performed on protein backbone atoms following structural alignment.
